# Presynaptic inhibition selectively suppresses leg proprioception in behaving *Drosophila*

**DOI:** 10.1101/2023.10.20.563322

**Authors:** Chris J. Dallmann, Yichen Luo, Sweta Agrawal, Grant M. Chou, Andrew Cook, Bingni W. Brunton, John C. Tuthill

**Affiliations:** Department of Physiology and Biophysics, University of Washington, Seattle, WA, USA; Department of Biology, University of Washington, Seattle, WA, USA; Department of Neurobiology and Genetics, Julius-Maximilians-University of Würzburg, Würzburg, Germany; School of Neuroscience, Virginia Tech, Blacksburg, VA, USA

**Keywords:** Motor control, proprioception, presynaptic inhibition, predictive signaling, efference copy, corollary discharge, ventral nerve cord, *Drosophila*

## Abstract

Controlling arms and legs requires feedback from proprioceptive sensory neurons that detect joint position and movement. Proprioceptive feedback must be tuned for different behavioral contexts, but the underlying circuit mechanisms remain poorly understood. Using calcium imaging in behaving *Drosophila*, we find that the axons of position-encoding leg proprioceptors are active across behaviors, whereas the axons of movement-encoding leg proprioceptors are suppressed during walking and grooming. Using connectomics, we identify a specific class of interneurons that provide GABAergic presynaptic inhibition to the axons of movement-encoding proprioceptors. The predominant synaptic inputs to these interneurons are descending neurons, suggesting they are driven by predictions of leg movement originating in the brain. Calcium imaging from both the interneurons and their descending inputs confirmed that their activity is correlated with self-generated but not passive leg movements. Overall, our findings elucidate a neural circuit for suppressing specific proprioceptive feedback signals during self-generated movements.

## Introduction

Effective motor control of arms and legs requires sensory feedback from proprioceptive sensory neurons (i.e., proprioceptors) that detect the position and movement of the body^1,2^. Motor circuits in the central nervous system integrate proprioceptive information at multiple levels to refine motor output and support a range of motor functions, from postural stabilization to adaptive locomotion^3–5^.

Because the same proprioceptors are used for many different motor control tasks, proprioceptive feedback must be flexibly tuned depending on the behavioral context^6^. For example, proprioceptive feedback pathways can be inhibited during voluntary movement to prevent disruption by reflexes^7,8^ or the inherent time delays in sensory pathways^9^. An efficient means of flexibly tuning sensory feedback pathways is predictive inhibition. In theoretical frameworks of predictive inhibition^10,11^, the motor circuits send an inhibitory signal to the sensory circuits that is based on the motor commands (Figure 1A). This mechanism, called efference copy or corollary discharge, allows self-generated sensory signals to be attenuated or eliminated, while externally-generated sensory signals are still transmitted to motor circuits.

**Figure 1.**
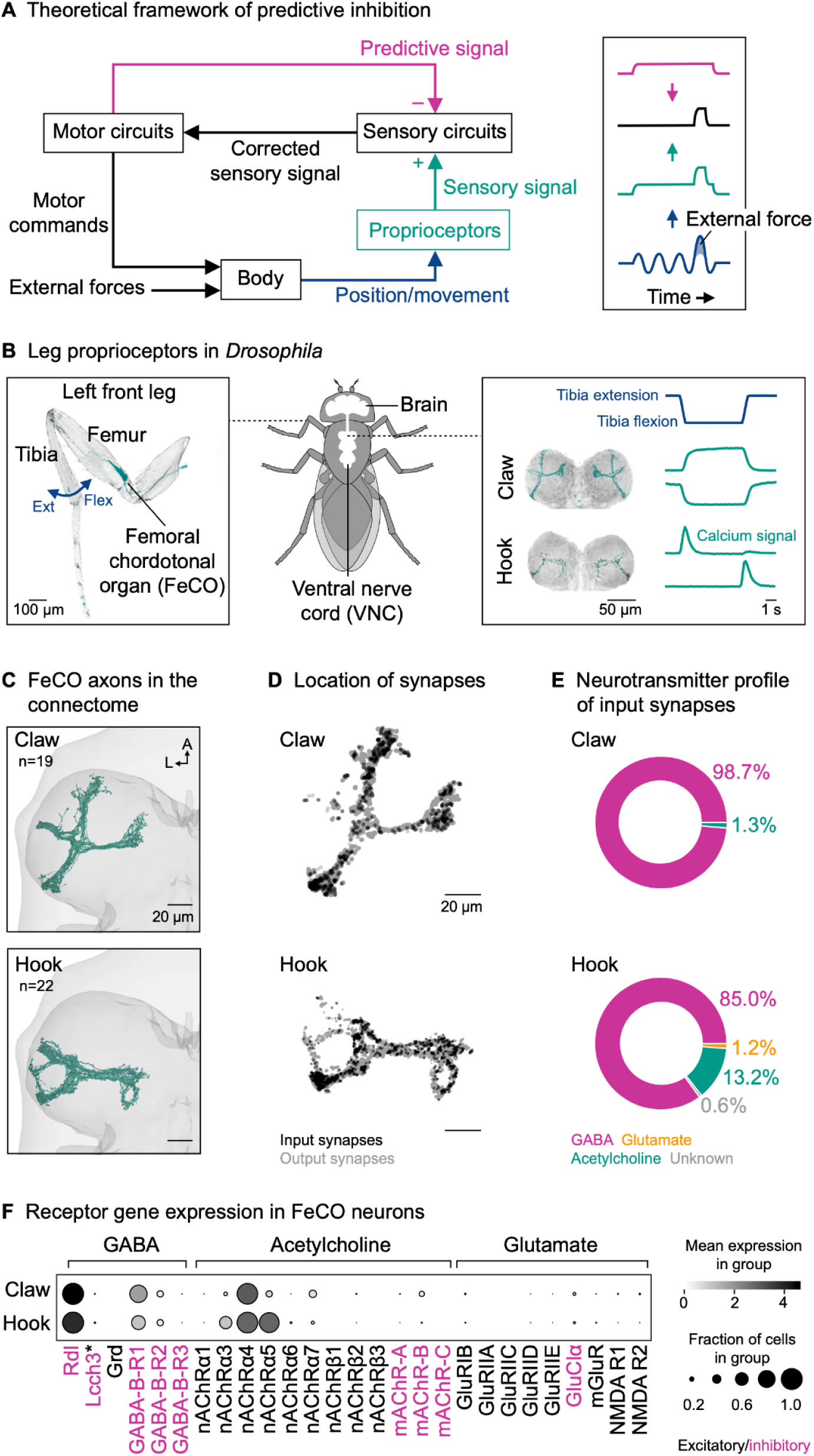
Proprioceptor axons from the *Drosophila* leg are positioned to receive presynaptic inhibition. (A) Left: Theoretical framework for predictive inhibition of proprioceptive pathways. Motor circuits send a predictive inhibitory signal (magenta) based on the motor commands to the sensory circuits. The predictive signal is subtracted from the measured sensory signal (green), representing a joint angle (blue) resulting from self-generated motor commands and external forces. The corrected sensory signal (black) can be used to counteract external forces without impeding voluntary movement. Right: Schematic time courses illustrating a situation where the predictive signal matches the sensory signal in timing and amplitude. (B) Left: Confocal image of a *Drosophila* front leg showing the location of the femoral chordotonal organ (FeCO) cell bodies and dendrites. Green: GFP; gray: cuticle auto-fluorescence. The blue arrow indicates the extension (Ext) and flexion (Flex) of the tibia relative to the femur. Right: Confocal image of position-encoding claw and movement-encoding hook axons in the fly ventral nerve cord (VNC). Flexion- and extension-encoding hook axons are overlaid. Green: GFP; gray: neuropil stain (nc82). Schematized calcium signals from claw and hook axons (GCaMP, green) in response to a controlled, passive movement of the femur-tibia joint (blue) based on Mamiya et al.^19^. (C) Top view of reconstructed claw and hook axons in the left front leg neuromere of the FANC connectome. n: number of axons; A: anterior; L: lateral. (D) Location of input and output synapses of all reconstructed claw and hook axons. View as in (C). (E) Neurotransmitter profile of the input synapses of claw and hook axons. (F) Expression levels of receptor genes in claw and hook neurons. Black intensity represents the mean level of gene expression in a cluster relative to the level in other clusters. Dot size represents the percent of cells in which gene expression was detected. Asterisk indicates that Lcch3 forms inhibitory channels with Rdl and excitatory channels with Grd^24^. See also Video S1.

Predictive inhibition of sensory feedback has been described for many different sensory modalities and species^6,10–13^. Inhibition can occur at multiple levels of the nervous system, but a common mechanism is presynaptic inhibition, where inhibitory neurons directly target the sensory axon terminals in the spinal cord or invertebrate ventral nerve cord (VNC) to reduce neurotransmitter release^14,15^. Previous studies have shown that presynaptic inhibition can dynamically suppress sensory transmission in proprioceptive axons, consistent with the theoretical framework of predictive inhibition. For example, the axons of leg proprioceptors in mice^8,9^ and locusts^16^ receive GABAergic inhibition during walking and reaching. However, the extent to which specific proprioceptive feedback pathways are inhibited during behavior and the organization and recruitment of the underlying neural circuits remains unknown. This is due in part to a lack of comprehensive connectivity analyses of proprioceptive circuits in the spinal cord and VNC, and the technical difficulty of recording from identified neurons in these circuits in behaving animals.

Here, we address these challenges in the fruit fly, *Drosophila melanogaster*. We focus on the femoral chordotonal organ (FeCO), the largest proprioceptive organ in the fly leg^17^ (Figure 1B, left). Proprioceptors in the FeCO are functionally analogous to vertebrate muscle spindles^2^. Genetically distinct “claw” and “hook” FeCO neurons monitor the position and movement of the tibia, respectively^18,19^. Position-encoding claw neurons are tonically active at different joint angles, whereas movement-encoding hook neurons are phasic and directionally tuned (Figure 1B, right). Feedback from FeCO neurons is integrated by circuits in the VNC to control leg posture and movement^20–22^. Previous studies in *Drosophila* have characterized the sensory signals of FeCO neurons during passive (i.e., externally-imposed) leg movements^18,19^. However, it remains unknown whether FeCO neurons receive presynaptic inhibition during active (i.e., self-generated) leg movements, and if so, which circuits mediate this inhibition.

Using cell type specific calcium imaging in behaving flies, we reveal that the movement-encoding hook axons but not the position-encoding claw axons are suppressed during walking and grooming. Using connectomics, we identify a specific class of GABAergic interneurons that provide the majority of inhibitory presynaptic input to the hook axons. These interneurons receive input from multiple descending neurons. Calcium imaging from both interneurons and descending neurons reveals a circuit by which the brain suppresses expected proprioceptive movement feedback during self-generated leg movements.

## Results

### Proprioceptor axons from the *Drosophila* leg are positioned to receive presynaptic inhibition

We first examined proprioceptor axons from the fly’s left front leg reconstructed from an electron microscopy volume of a female *Drosophila* VNC (FANC^23^; Figure 1C; Video S1; Table S1).To determine whether proprioceptors receive presynaptic input, we analyzed the location and number of the input and output synapses of FeCO axons. Input synapses were present on all axon branches, spatially intermingled with output synapses (Figure 1D). On average, individual claw and hook axons had 29 ± 18 (mean ± std) and 63 ± 41 input synapses and 744 ± 311 and 787 ± 380 output synapses, respectively. Individual axons differed in the total number of synapses; the number of input synapses was positively correlated with the number of output synapses (claw: r(17) = 0.79, p < 0.001; hook: r(20) = 0.45, p = 0.035). By identifying the neurons presynaptic to claw and hook axons (see STAR methods), we found that the presynaptic partners were primarily GABAergic (Figure 1E). Consistent with this finding, we analyzed a single-cell RNA-sequencing dataset^19^ and found that all claw and hook neurons strongly express Rdl, the gene for GABA_A_ receptors (Figure 1F). In contrast, the gene for the inhibitory glutamate receptor (GluClα) was only weakly expressed in a few FeCO cells (Figure 1F).

Together, connectomic reconstruction and analysis of gene expression from claw and hook neurons suggest that they receive GABAergic input from interneurons in the VNC, which provides a substrate for context-dependent presynaptic inhibition.

### Tools to study leg proprioception in behaving *Drosophila*

To investigate the function of presynaptic inhibition of FeCO axons, we developed a setup for two-photon calcium imaging of neural activity in the VNC and 3D leg tracking of tethered flies on an air-supported ball, which functions as an omnidirectional treadmill (Figure 2A; see STAR methods). The setup allowed us to record calcium signals in FeCO axons and other neurons in the VNC with the genetically-encoded calcium sensor GCaMP while flies walked, groomed, or rested on the treadmill.

**Figure 2.**
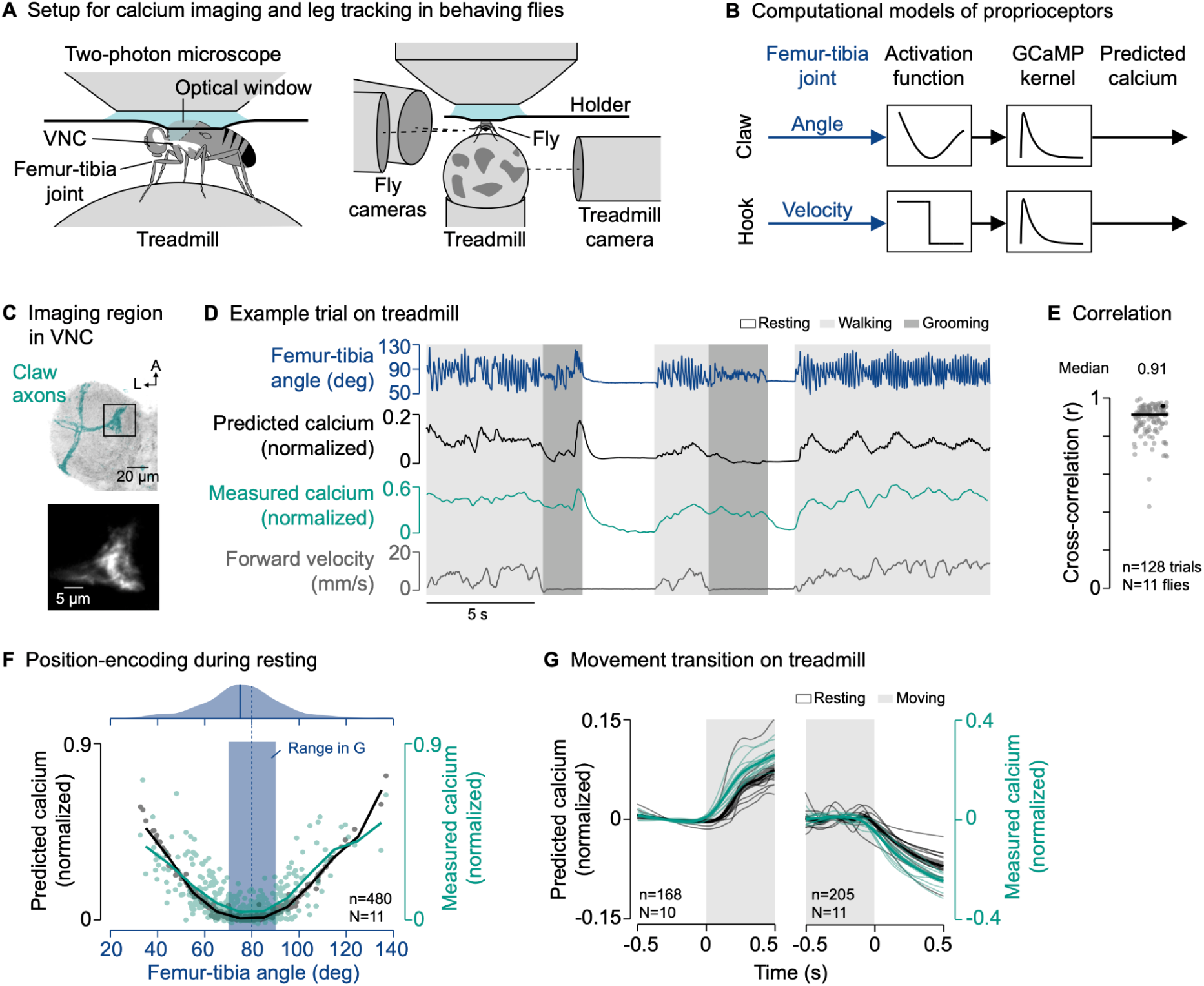
The axons of position-encoding proprioceptors are not suppressed during active leg movements. (A) Experimental setup for two-photon calcium imaging from VNC neurons and 3D leg tracking of the left front leg of tethered flies on a treadmill. (B) Computational models of FeCO proprioceptors translating time courses of joint angles into time courses of calcium signals. The activation functions were fitted to calcium signals measured during passive leg movement. (C) Top: Confocal image of position-encoding claw axons in the VNC. The black box indicates the imaging region. Green: GFP; gray: neuropil stain (nc82). A: anterior; L: lateral. Bottom: Mean tdTomato signal within the imaging region during an example trial. (D) Example trial of two-photon calcium imaging of claw axons in the neuromere of the left front leg and behavior tracking on the treadmill. (E) Cross-correlation coefficient between predicted and measured calcium signals per trial at a time lag of zero. The black line shows the median. The black dot marks the trial shown in (D). n: number of trials; N: number of flies. (F) Median predicted and measured calcium signals as a function of the median femur-tibia angle for individual resting bouts. Bouts are ≥1 s in duration. The black and green line indicate the mean calcium signals in bins of 10°. The dashed blue line indicates the resting angle at which activity is minimal. The blue rectangle indicates the range of resting angles analyzed in (G). The plot on top shows a kernel density estimation of the femur-tibia angles during resting. The solid blue line indicates the most frequent femur-tibia angle (mode of the distribution). n: number of resting bouts; N: number of flies. (G) Predicted and measured calcium signals aligned to the transitions into and out of movement. Movement includes walking and grooming. Thin lines show animal means, thick lines show mean of means, shadings show standard error of the mean. n: number of transitions; N: number of flies. See also Figures S1 and S2 and Video S2.

To compare the neural recordings with hypotheses about circuit function, we constructed computational models that predicted calcium signals in the neurons of interest based on behavior (see STAR Methods). The models convolved a neuron-specific activation function with a GCaMP kernel to translate time courses of joint kinematics or binary behavioral signals into time courses of calcium signals (Figure 2B).

In the case of claw and hook neurons, the activation function within each model was based on our previous calcium imaging and leg tracking data, in which the femur-tibia joint was passively moved^18^ (Figure S2A). As a population, claw neurons encode the position of the femur-tibia joint as a deviation from a joint angle of ∼80°. Population activity increases non-linearly with increasing flexion or extension (Figure S2B). In contrast, hook neurons respond transiently to flexion or extension of the femur-tibia joint (Figure S2D). Our claw and hook models effectively replicated these characteristic calcium signals during passive leg movements (Figure S2B and S2D). This was reflected in high cross-correlation coefficients between measured and predicted calcium signals across trials and flies (claw: r = 0.93; hook: r = 0.85; Figure S2C and S2E). We then used these models to predict calcium signals in FeCO axons during active leg movements (Figure 2D). This comparison of predicted calcium signals based on passive leg movements and measured calcium signals during active leg movements provided a quantitative means to identify context-dependent inhibition. The comparison was particularly useful for interpreting calcium signals in claw axons, whose position-sensitivity led to unintuitively weak signals in some movement bouts that could be mistaken for inhibition without a model prediction (e.g., Figure S2G).

### The axons of position-encoding proprioceptors are not suppressed during active leg movements

Equipped with computational models to predict calcium signals during active leg movements, we first asked whether the axons of the position-encoding claw neurons are suppressed during behavior. We co-expressed the calcium indicator GCaMP and the structural marker tdTomato (for motion correction) with the same cell-specific genetic driver line^18^ that was used to tune the passive computational model (Figure 2C and S1), and recorded the activity of claw axons in behaving flies.

Claw axons were active across behaviors—during resting, walking, and grooming (Figure 2D and S2F; Video S2). The passive model effectively tracked the temporal dynamics of the calcium signal across different behaviors (Figure 2D). This was reflected in high cross-correlation coefficients between measured and predicted calcium signals across trials and flies (r = 0.91; Figure 2E), which were comparable to those in the passive movement experiments used to tune the passive model (Figure S2C). Calcium signals were also well predicted when we removed the treadmill and flies moved their legs freely in the air (Figure S2G and S2H; Video S2).

The characteristic position-encoding of claw axons was particularly clear when resting flies held their front leg at a given femur-tibia angle for an extended period of time. Plotting the median amplitude of the calcium signal against the median femur-tibia joint angle for individual resting bouts (≥1 s in duration) revealed the expected U-shaped activity pattern centered at ∼80° (Figure 2F, bottom). The minimum signal was close to the most frequent femur-tibia angle that flies adopted while resting on the treadmill (75°; Figure 2F, top).

Given this U-shaped activity pattern, we expected to see strong changes in the calcium signal when flies transitioned between resting and moving near the most frequent resting angle. Indeed, for transitions toward or away from resting angles of 70°-90° (Figure 2F, blue box), calcium signals increased and decreased as predicted by the passive model (Figure 2G).

Together, these results indicate that the position-encoding claw axons are not suppressed during self-generated leg movements, and that position feedback is transmitted to downstream VNC neurons across behavioral contexts. The close match between the passive claw model predictions and calcium signals recorded during behavior also provides confidence in our approach of comparing self-generated and passive leg movements in leg proprioceptors.

### The axons of movement-encoding proprioceptors are suppressed during active leg movements

We next asked whether the axons of movement-encoding proprioceptors are suppressed during behavior. We first investigated hook neurons encoding tibia flexion movements. We again co-expressed the calcium indicator GCaMP and the structural marker tdTomato in the same cell-specific driver line^18^ used for tuning the passive computational model (Figure 3A and S1), and recorded the activity of hook axons in behaving flies.

**Figure 3.**
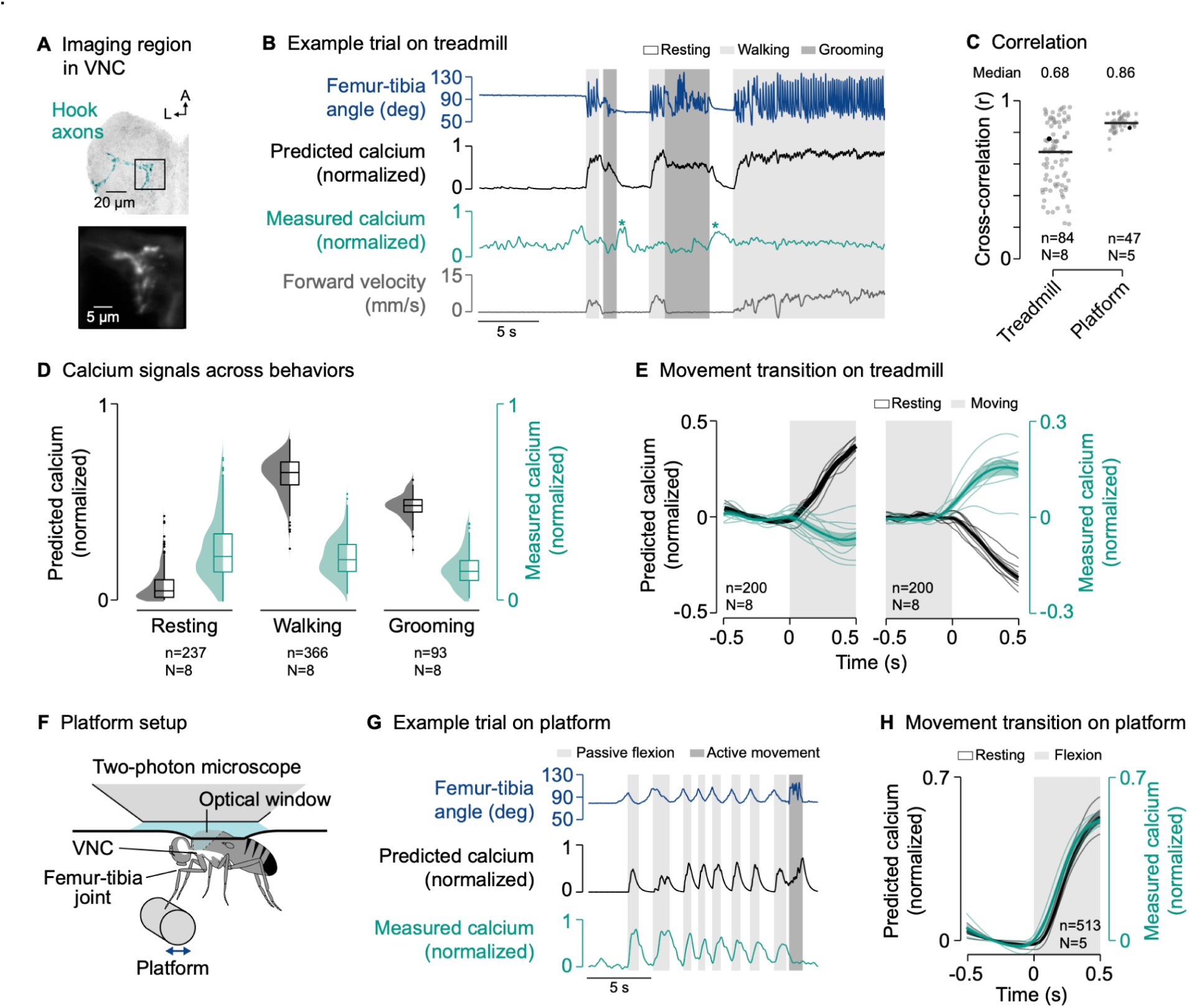
The axons of movement-encoding proprioceptors are suppressed during active leg movements. (A) Top: Confocal image of movement-encoding hook (flexion) axons in the VNC. The black box indicates the imaging region. Green: GFP; gray: neuropil stain (nc82). A: anterior; L: lateral. Bottom: Mean tdTomato signal within the imaging region during an example trial. (B) Example trial of two-photon calcium imaging of hook flexion axons in the neuromere of the left front leg and behavior tracking on the treadmill. The asterisks highlight resting bouts during which the front leg was held in the air and slowly flexed, likely as a result of passive forces produced by leg muscles and skeletal structures. (C) Cross-correlation coefficient between predicted and measured calcium signals per trial at a time lag of zero in different movement contexts. Black lines show medians. Black dots mark the trials shown in (B) and (G). In platform trials, active movements were excluded for the cross-correlation. n: number of trials; N: number of flies. (D) Median predicted and measured calcium signals during resting, walking, and grooming. Bouts are ≥1 s in duration. Distributions show kernel density estimations. n: number of behavioral bouts; N: number of flies. (E) Predicted and measured calcium signals aligned to the transitions into and out of movement. Signals are baseline subtracted (mean from −0.5 to 0 s). Movement includes walking and grooming. Thin lines show animal means, thick lines show mean of means, shadings show standard error of the mean. n: number of transitions; N: number of flies. (F) Experimental setup for passively moving the left front leg via a platform during two-photon calcium imaging from the VNC. (G) Example trial of two-photon calcium imaging of hook flexion axons and behavior tracking on the platform. (H) Predicted and measured calcium signals aligned to the transition into passive flexion of the femur-tibia joint. Lines and labels as in (E). See also Figures S1 and S3 and S4 and S5 and S6 and Video S3.

The passive model predicted strong calcium signals in hook axons during walking and grooming compared to resting (Figure 3B and 3D). However, hook calcium signals recorded during behavior were conspicuously different from the passive model predictions (Figure 3B and 3D; Video S3). The discrepancy between model prediction and measurement was particularly obvious at the transitions into and out of movement, at which calcium signals did not increase or decrease as predicted by the passive model (Figure 3E). Accordingly, the cross-correlation between measured and predicted calcium signals across trials and flies was more variable and lower on average than during passive movement (r = 0.68; Figure 3C, Treadmill). Note that we computed high cross-correlation coefficients in some trials simply because those flies did not frequently transition between different behaviors. Calcium signals were also absent when we removed the treadmill and flies moved their legs freely in the air (Figure S3A-C; Video S3). The lack of calcium signals during self-generated movements was even more pronounced in a second driver line^19^ for hook flexion neurons (Figure S3D-I). We also observed a similar degree of context-dependent suppression in hook neurons encoding tibia extension^19^ movements (Figure S4). These results indicate that both flexion- and extension-encoding hook axons are suppressed during self-generated but not passive leg movements.

Supporting this conclusion, we observed that calcium signals were high when the front leg was moved passively while resting on the treadmill, which sometimes occurred when the hind legs lifted off the treadmill for grooming (Video S3). Calcium signals were also high during resting when the front leg slowly (over the course of hundreds of milliseconds) moved towards flexion, which we observed after front leg grooming when the leg was not on the treadmill (Figure 3B, asterisks) or in trials in which the treadmill was removed (Figure S3A). These slow flexions were likely the result of passive forces produced by leg muscles^25^ and skeletal structures^26^.

To further test that hook axons are not suppressed during passive movements, we replaced the treadmill with a moveable platform that flies gripped with the tips of their legs (Figure 3F; Video S3). We used the platform to passively move the front leg while imaging from hook axons in the VNC. In this context, we measured strong calcium signals in response to passive movement of the femur-tibia joint, as predicted by the passive model (Figure 3G and 3H). This was reflected in higher and less variable cross-correlation coefficients between predicted and measured calcium signals across trials and flies (r = 0.86; Figure 3C, Platform). Because flies were not anesthetized, they sometimes actively moved their legs instead of gripping the platform. In line with our previous findings, calcium signals were weak during these active movements (Figure 3G and S3J).

Finally, we asked whether differences between the passive model predictions and recorded calcium signals could be due to differences in joint movement dynamics between the active and passive movement conditions. Specifically, flies tended to move their legs more rapidly when they were actively moving compared to how we moved them during passive stimulation with the platform. Using the same magnetic control system in which we previously investigated proprioceptor responses to passive movements^18,19^, we replayed naturalistic time courses of femur-tibia joint angles measured during walking and grooming to otherwise passive animals (Figure S5A and S5B; see STAR methods). Calcium signals recorded from hook axons in this passive context matched the predictions of the passive model, with calcium signals increasing at the onset of movement as predicted (Figure S5C-H). Thus, the discrepancy between activity recorded during self-generated movements and the passive model predictions is unlikely to be caused by differences in stimulus statistics.

Together, these results indicate that movement-encoding hook axons are suppressed whenever flies move their legs actively, regardless of the specific movement context.

In addition to claw and hook neurons, the FeCO contains a third class of neurons (“club”), which encode low-amplitude, high-frequency vibrations^18,19^ and are thought to function as exteroceptors rather than proprioceptors^20,21^. We used a cell-specific driver line^19^ to record the calcium activity in the axons of club neurons and found that they are not suppressed during active leg movements (Figure S6A-G and S1; Video S4), similar to claw axons. Interestingly, the baseline activity in these axons was elevated when the legs contacted the treadmill (Figure S6H-I), consistent with the idea that the neurons function as exteroceptors that mediate external substrate vibrations.

### GABAergic interneurons provide presynaptic inhibition to movement-encoding proprioceptor axons

To explore the circuit mechanisms underlying the selective suppression of hook axons, we analyzed the presynaptic connectivity of claw and hook axons in the female VNC connectome (Figure 4A). Claw and hook axons receive synaptic input from interneurons and sensory neurons intrinsic to the VNC, but not descending neurons (claw: 100% of input synapses from interneurons; hook: 94.9% interneurons, 4.6% sensory neurons, 0.5% unknown). On average, hook axons receive more presynaptic input than claw axons (Figure 4A, top), with most input coming from GABAergic interneurons (Figure 4A, left). Interestingly, presynaptic neurons target either claw axons or hook axons, but not both. This could explain why activity was selectively suppressed in hook axons but not claw axons. Most GABAergic input onto hook axons (83.1% of input synapses) comes from a group of local interneurons belonging to the 9A hemilineage^27,28^ (Table S1). One 9A neuron in particular provides 56.8% of presynaptic input to hook axons (Figure 4A, right). This chief 9A neuron receives dendritic input in the dorsal VNC and provides synaptic output to hook axons in the ventral VNC (Figure 4B; Video S1). In fact, most of the output of the chief 9A neuron (62.8%) is onto hook axons (Figure 4C). Other GABAergic neurons of the 9A hemilineage also provide a significant fraction of their output to hook axons (Figure 4C). Thus, this group of GABAergic 9A interneurons is positioned to selectively suppress activity in hook axons during active leg movements via presynaptic inhibition.

**Figure 4.**
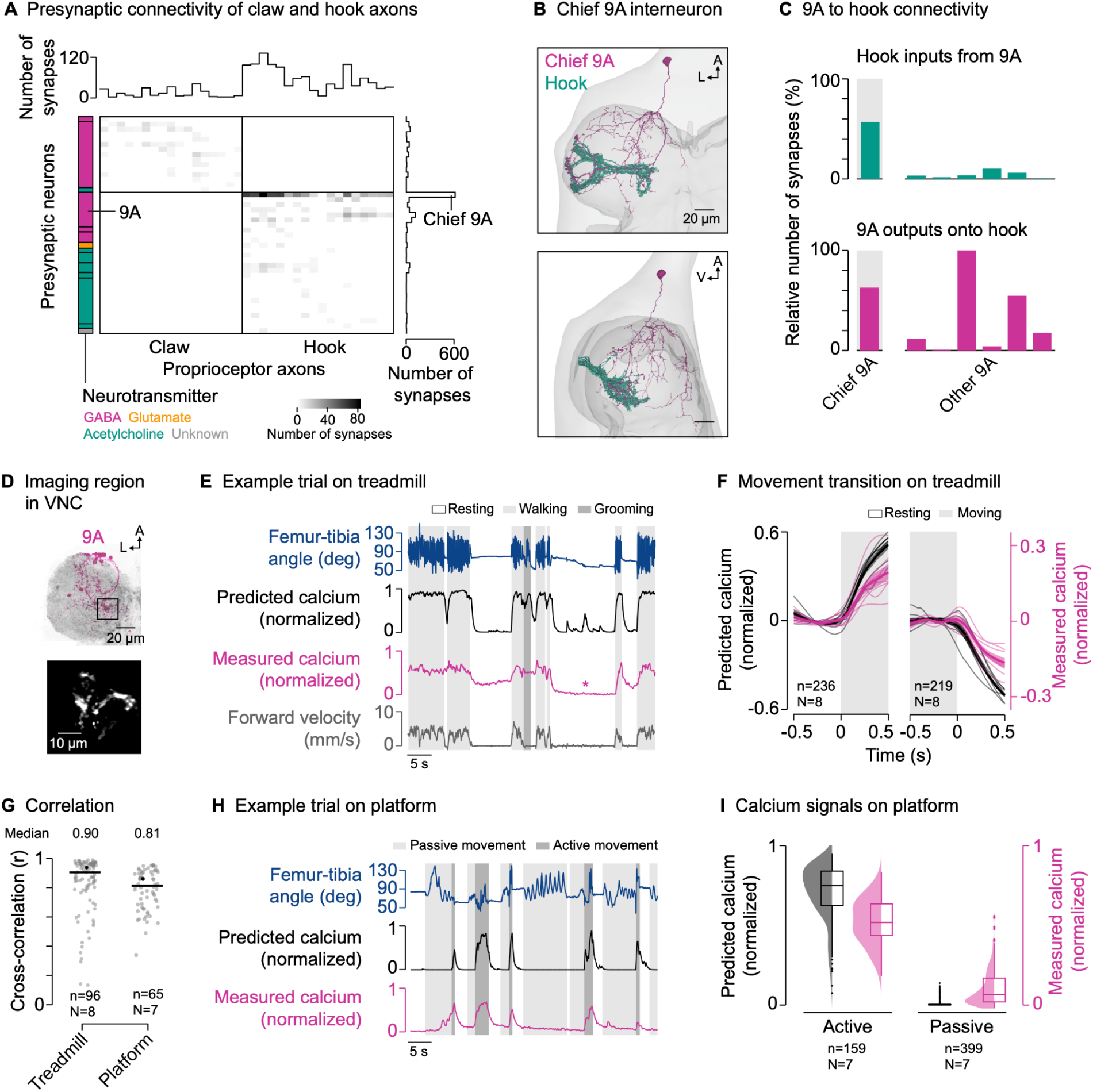
GABAergic interneurons provide presynaptic inhibition to movement-encoding proprioceptor axons. (A) Connectivity of presynaptic neurons with claw and hook axons. The grayscale heatmap indicates the number of synapses between neurons (connection strength). Boxes on the left group presynaptic neurons of the same developmental lineage, with the color indicating their primary fast-acting neurotransmitter. Boxes from top to bottom: 13B and 19A (both GABA); 3A (acetylcholine); 9A, 13B, and 19A (all GABA); 8A (glutamate); 1A, 8B, 18B, 22A, hook axons, and hair plate axon (all acetylcholine); unknown. (B) Top and side view of the chief GABAergic 9A neuron presynaptic to hook axons in the left front leg neuromere in FANC. A: anterior; L: lateral; V: ventral. (C) Connectivity between 9A neurons and hook axons. (D) Top: Confocal image of 9A neurons in the VNC. The black box indicates the imaging region. Magenta: GFP; gray: neuropil stain (nc82). A: anterior; L: lateral. Bottom: Mean tdTomato signal within the imaging region during an example trial. (E) Example trial of two-photon calcium imaging of 9A neurons in the neuromere of the left front leg and behavior tracking on the treadmill. The asterisk highlights a resting bout during which the front leg was moved passively by the grooming hind legs. (F) Predicted and measured calcium signals aligned to the transitions into and out of movement. Movement includes walking and grooming. Thin lines show animal means, thick lines show mean of means, shadings show standard error of the mean. n: number of transitions; N: number of flies. (G) Cross-correlation coefficient between predicted and measured calcium signals per trial at a time lag of zero in different movement contexts. Black lines show medians. Black dots mark the trials shown in (E) and (H). n: number of trials; N: number of flies. (H) Example trial of two-photon calcium imaging of 9A neurons and behavior tracking on the platform. (I) Median predicted and measured calcium signals during active and passive movement bouts on the platform. Bouts are ≥1 s in duration. Distributions show kernel density estimations. n: number of movement bouts; N: number of flies. See also Figure S7 and Videos S1 and S5.

If these 9A neurons suppress activity in hook axons, we would expect their activity to be high during active leg movements and low during passive leg movements; the opposite activity pattern that we observed in hook axons. We instantiated this prediction in a simple computational model, in which calcium activity is high during active flexion and extension movements, but not during resting or passive leg movements (Figure S2A; see STAR methods). We then tested the predictions of the model by recording from axons of 9A neurons in a region near the terminals of hook axons in the front leg neuromere using a cell-specific genetic driver line (Figure 4D and S1). As predicted, we measured strong calcium signals in the axons of the 9A neurons during walking and grooming (Figure 4E; Video S5), with calcium signals increasing and decreasing at the transitions into and out of movement, respectively (Figure 4F). This was reflected in high cross-correlation coefficients between predicted and measured calcium signals across trials and flies (r = 0.90; Figure 4G). Calcium signals were also well predicted when we removed the treadmill and flies moved their legs freely in the air (r = 0.95; Figure S7A-C; Video S5). Notably, calcium signals in 9A neurons were weak when the front leg was at rest while other legs moved actively, as was the case during hind leg grooming (Figure S7D). This suggests that 9A neurons can be recruited in a leg-specific manner.

Calcium signals in 9A neurons were also weak when the grooming hind legs passively moved the front leg (Figure 4E, asterisk; Video S5). To further test whether calcium signals in 9A axons are absent during passive leg movements, we again used the platform setup to passively move the femur-tibia joint (Figure 4H; Video S5). Because flies were not anesthetized, they sometimes actively moved their legs instead of gripping the platform. As predicted by the computational model, calcium signals were weak during passive leg movements and strong during active leg movements (Figure 4H and 4I), with high cross-correlation coefficients between predicted and measured calcium signals (r = 0.81; Figure 4G).

Together, these results demonstrate that local GABAergic 9A neurons are active during self-generated but not passive leg movements. The synaptic connectivity and activity pattern of these neurons suggest that they selectively suppress hook axons via presynaptic inhibition.

To test the behavioral significance of the 9A neurons, we used optogenetics to transiently (1 s) activate or silence the neurons in tethered flies walking on a treadmill. We expressed CsChrimson^29^ or GtACR1^30,31^ in 9A neurons and used a red or green laser focused on the ventral thorax at the base of the left front leg^20,32^ to specifically manipulate the 9A neurons in the neuromere of that leg (Figure S7E; see STAR methods). Flies were able to walk during the manipulations (Figure S7F). Activating 9A neurons changed the range of femur-tibia movements and caused flies to slow down (Figure S7G-H, top). Silencing 9A neurons had weaker effects (Figure S7G-H, bottom). Overall, these results are consistent with a role for 9A neurons in suppressing leg proprioceptors that contribute to local feedback control of the femur-tibia joint. They are also consistent with past work showing that flies lacking feedback from FeCO proprioceptors can still walk on even terrain, though they exhibit subtly altered step kinematics^22,33,34^.

### GABAergic interneurons receive descending input from the brain

To explore the origins of context-dependent activity in the GABAergic 9A neurons, we analyzed their presynaptic neurons in the connectome (Figure 5A). The 9A neurons receive little direct input from sensory neurons, suggesting they are not driven by sensory feedback from the leg. Interestingly, the chief 9A neuron, which provides the majority of the input to hook axons, receives most of its input (68.3% of input synapses) from descending neurons (Table S1). The other 9A neurons receive most of their input from local premotor neurons in the VNC, but some neurons of the group also receive substantial descending input (Figure 5B). In fact, several specific descending neurons provide input to multiple 9A neurons (Figure 5B and S8A). These descending neurons have no or only few synapses with the GABAergic neurons presynaptic to claw axons (Figure S8A). Thus, the 9A neurons may be recruited together by descending input from the brain to presynaptically inhibit hook axons during behavior.

**Figure 5.**
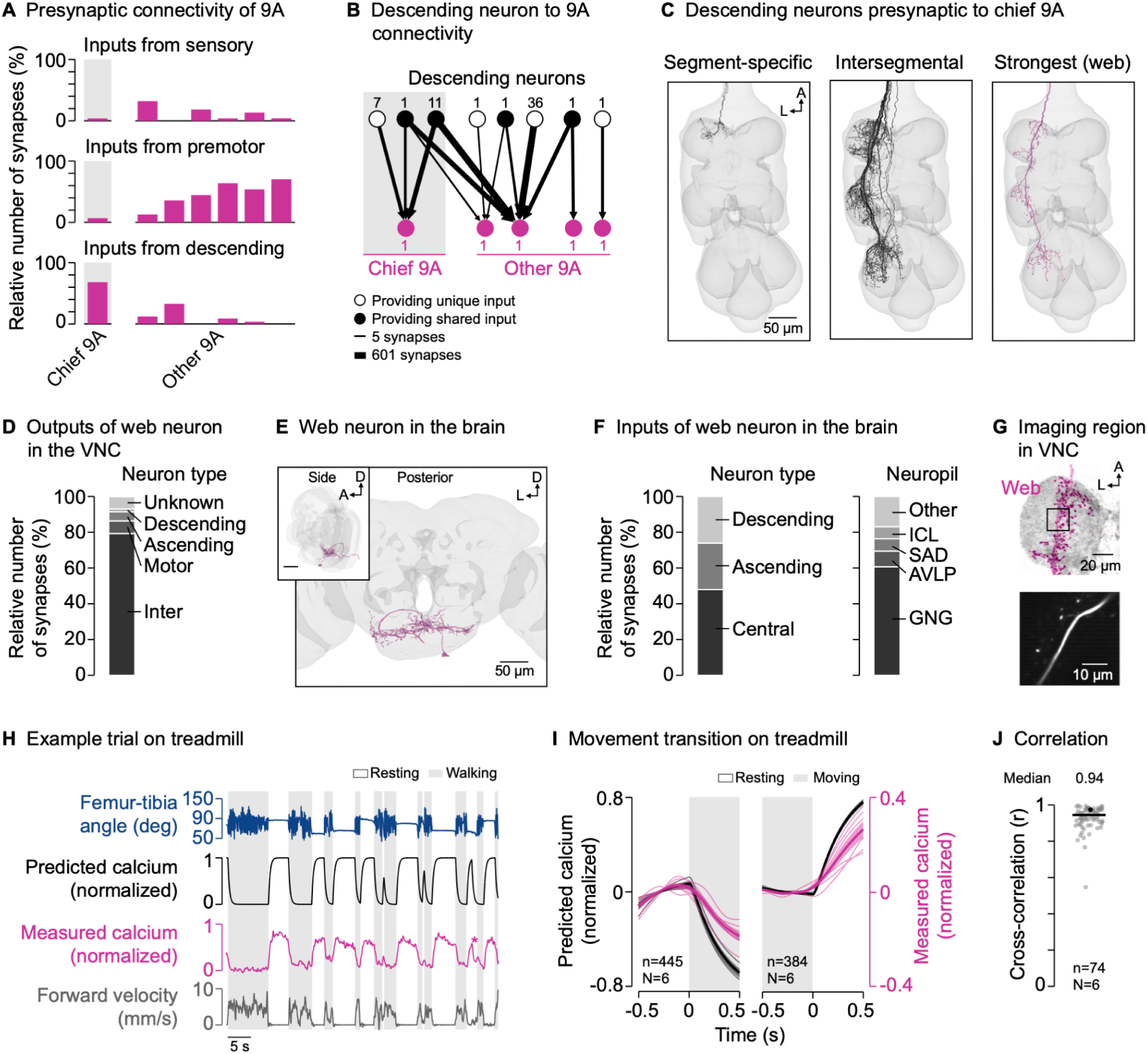
GABAergic interneurons receive descending input from the brain. (A) Inputs from sensory neurons, premotor neurons, and descending neurons onto individual 9A neurons presynaptic to hook axons in the front leg neuromere (FANC connectome). (B) Connectivity between descending neurons and the 9A neurons presynaptic to hook axons (FANC connectome). Numbers next to nodes indicate the number of neurons with the same connectivity motif. Lines indicate the log10 of the number of synapses. (C) Segment-specific and intersegmental descending neurons presynaptic to chief 9A, including the most strongly connected descending neuron (web; FANC connectome). A: anterior; L: lateral. (D) Outputs of the descending web neuron in the VNC (MANC connectome). (E) Posterior and side view of the descending web neuron in the brain (FlyWire connectome). A: anterior; D: dorsal; L: lateral. (F) Inputs to the descending web neuron in the brain (FlyWire connectome). GNG: gnathal ganglia; AVLP: anterior ventrolateral protocerebrum; SAD: saddle; ICL: inferior clamp. (G) Top: Confocal image of web neuron in the VNC. The black box indicates the imaging region. Magenta: GFP; gray: neuropil stain (nc82). A: anterior; L: lateral. Bottom: Mean tdTomato signal within the imaging region during an example trial. (H) Example trial of two-photon calcium imaging of the web neuron in the neuromere of the left front leg and behavior tracking on the treadmill. The asterisk highlights part of a front leg resting bout during which the hind legs were grooming. (I) Predicted and measured calcium signals aligned to the transitions into and out of movement. Movement includes walking and grooming. Thin lines show animal means, thick lines show mean of means, shadings show standard error of the mean. n: number of transitions; N: number of flies. (J) Cross-correlation coefficient between predicted and measured calcium signals per trial at a time lag of zero. The black line shows the median. The black dot marks the trial shown in (H). n: number of trials; N: number of flies. See also Figure S8 and Videos S1 and S6.

Some of the descending neurons presynaptic to the chief 9A neuron target only the neuromere of the left front leg (Figure 5C, left), confirming our observation that 9A neurons can be controlled in a leg-specific manner. However, the majority of descending input (76.1%) comes from intersegmental descending neurons that target multiple leg neuromeres of one body side (Figure 5C, middle). This raised the possibility that the circuit motif we identified for the left front leg is present in all legs. To test this possibility, we turned to a second VNC connectome of a male fly, which is more fully reconstructed in the middle and rear neuromeres (MANC^35,36^; see STAR methods). In support of our hypothesis, we found that hook axons in all leg neuromeres in the male connectome receive most of their input from a 9A neuron resembling the chief 9A neuron in the female connectome (Figure S8B; Table S2). These chief 9A neurons target primarily hook axons (58.0% of output synapses on average) and provide little output to other sensory axons (12.9% of output synapses on average; Figure S8D). Moreover, the top descending neurons presynaptic to the chief 9A neurons resemble those in the female connectome (Figure S8C; Table S2). Thus, the inhibitory circuit motif is present in both females and males and segmentally repeated to primarily inhibit hook axons from all legs.

In the female VNC connectome, some of the descending neurons presynaptic to chief 9A were recently annotated (Table S1 and Figure S8A). Three of these descending neurons are intersegmental cholinergic neurons that drive walking^37^ (BDN2, oDN1) and turning^38^ (DNa02), two of which (BDN2 and DNa02) were shown to be normally active during walking^37,38^. Another annotated descending neuron is a cholinergic neuron that drives front leg grooming^39^ (DNg12). This suggests that excitatory descending neurons can recruit the chief 9A neuron to drive feedback inhibition during walking and grooming, in parallel to acting on motor circuits in the VNC.

The descending neuron providing the most input to chief 9A had not been annotated in the female VNC connectome. However, its match in the male VNC connectome was recently annotated^40^ as a GABAergic “web” neuron, a type of descending neuron targeting primarily the leg neuropils^41^ (Figure 5C, right; Table S2). We found that, in the VNC, the web neuron targets primarily interneurons (79.4% of output synapses), including the chief 9As (1.9%; Figure 5D). To explore which brain regions provide input to the web neuron, we turned to a brain connectome of a female fly (FlyWire^42–44^; Figure 5E; Table S3; see STAR methods). We found that the web neuron receives input from neurons central to the brain (59 neurons, 48.0% of input synapses), ascending neurons (74 neurons, 26.6%), and descending neurons (37 neurons; 25.3%; Figure 5F, left). Most of the brain input (60.7%) comes from the gnathal ganglia (GNG), a brainstem-like region important for descending locomotor control^45^ (Figure 5F, right).

If the GABAergic web neuron helps recruit the chief 9A neuron during active leg movements, we would expect its activity to be low during active leg movements (corresponding to disinhibition) and high during passive leg movements or resting; the opposite activity pattern that we observed in the 9A neurons. We instantiated this prediction in a simple computational model, in which calcium activity is high during resting but low during active leg movements (see STAR methods). We then tested the predictions of the model by recording from the web neuron in the front leg neuromere using a cell-specific driver line^41^ (Figure 5G and S1). As predicted, we measured strong calcium signals during resting but not during walking (Figure 5H and Video S6), with calcium signals decreasing and increasing at the transitions into and out of movement, respectively (Figure 5I). This was reflected in high cross-correlation coefficients between predicted and measured calcium signals across trials and flies (r = 0.94; Figure 5J). Calcium signals were also weak when we removed the treadmill and flies moved their legs freely in the air (Figure S8E; Video S6). Notably, calcium signals in the web neuron were also weak when the front leg was at rest while other legs moved actively, as was the case during hind leg grooming (Figure 5H, asterisk). This suggests that the GABAergic web neuron disinhibits its postsynaptic target neurons throughout the VNC during self-generated leg movements.

Together, these results support a circuit motif in which excitatory and inhibitory descending signals from motor circuits in the brain drive the 9A neurons in a context-dependent manner (Figure 6A). During active leg movements like walking and grooming, excitatory descending neurons recruit the 9A neurons in parallel to acting on motor circuits in the VNC, while inhibitory descending neurons release them from inhibition (Figure 6B, left). During passive leg movements or resting, 9A neurons are not recruited due to the missing excitatory drive and the inhibition by descending neurons (Figure 6B, right). The identified excitatory descending neurons that drive walking are intersegmental; they are positioned to recruit 9A neurons in all leg neuromeres (Figure 6C, left). The excitatory descending neurons that drive front leg grooming are segment-specific; they are positioned to recruit 9A neurons for the front legs, leaving proprioceptive transmission in the standing middle and hind legs unaffected (Figure 6C, right).

**Figure 6.**
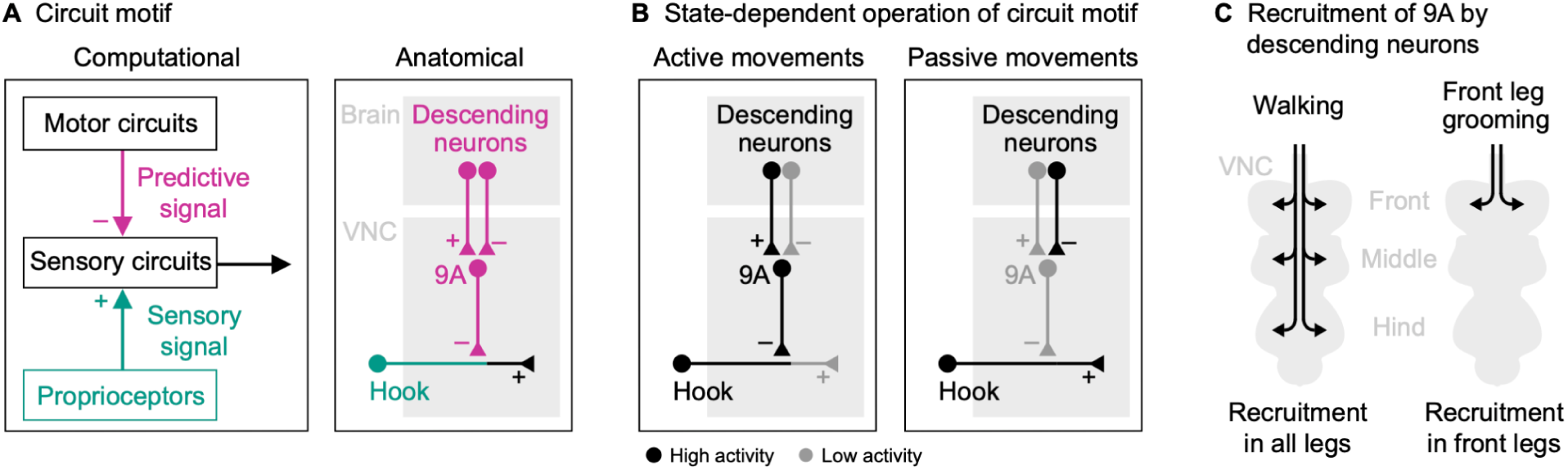
Summary of neural circuit for selectively suppressing proprioceptive movement feedback. (A) A set of descending neurons target GABAergic 9A neurons in the VNC, which suppress feedback from movement-encoding hook axons via presynaptic inhibition during active, self-generated leg movements. (B) Context-dependent operation of the circuit motif during active leg movements and passive leg movements/resting. (C) Recruitment of 9A neurons in different VNC neuropils by excitatory descending neurons that drive walking and front leg grooming.

## Discussion

In this study, we elucidate a neural circuit that selectively suppresses proprioceptive feedback from the *Drosophila* leg in a context-dependent manner. Previous studies have shown that proprioceptive pathways are modulated by inhibition during self-generated leg movements^8,9,16^. However, how specific pathways are inhibited during behavior and the organization and recruitment of the underlying neural circuits was previously unknown. We leveraged connectomics and neural recordings in behaving animals to show that the movement-encoding hook but not the position-encoding claw axons from the femoral chordotonal organ (FeCO) are suppressed during self-generated leg movements (Figures 2 and 3). The hook axons receive GABAergic presynaptic inhibition from a specific class of interneurons, which are exclusively active during self-generated leg movements (Figure 4). These neurons receive input from descending neurons in the brain, whose activity suggests that they drive feedback inhibition in a predictive manner (Figures 5 and 6).

### GABA-mediated suppression of hook axons

Our findings suggest that hook axons are suppressed via GABAergic presynaptic inhibition. The connectome revealed overwhelmingly GABAergic input onto hook axons, and RNA-seq revealed that hook neurons strongly express the GABA_A_ receptor gene Rdl. GABAergic presynaptic inhibition of somatosensory axons is common throughout the animal kingdom^15^. For example, the axons of leg proprioceptors of locusts^16^ and mice^8^ receive GABAergic presynaptic inhibition during walking. Notably, presynaptic inhibition can also be mediated by other neurotransmitters, including glutamate^15,46^. In the case of hook axons, however, the low number of glutamatergic input synapses in the connectome and the low expression level of the inhibitory glutamate receptor gene GluClα argue against glutamatergic inhibition as the driving force.

### Behavioral function of sensory suppression in hook axons

We found that the movement-encoding hook axons were suppressed during all self-generated leg movements. Attenuating expected proprioceptive feedback could increase the animal’s sensitivity to external perturbations and facilitate compensatory motor actions. An advantage of movement over position feedback in this context is that perturbations can be detected more rapidly. This mechanism would require the transmission of externally-generated signals during movement. The high speed of leg movement during fly walking and the slow dynamics of the calcium signal prevented us from testing this hypothesis directly (see below).

Inhibition of proprioceptive feedback can also function to attenuate reflexes that would disrupt ongoing movements^7^. This is seen in mice, where chronic removal of a large population of inhibitory interneurons in the spinal cord causes excessive leg flexion that disrupts walking^8^. In big insects, neurons of the FeCO are known to mediate postural reflexes that stabilize the leg against external perturbations^47–49^. In *Drosophila*, similar postural reflexes exist that could be mediated by FeCO neurons^32^. Consistent with this idea, we measured strong calcium signals in hook axons when the standing front leg was perturbed on the treadmill (for example during hind leg grooming; Video S3). The suppression of hook axons during active leg movements could then function to prevent the recruitment of a postural reflex. We found that transient silencing of the GABAergic 9A neurons had only subtle effects on walking. This suggests that feedback from hook axons alone is insufficient to recruit a disrupting reflex. We believe this is because the reflex is mediated by both movement and position feedback from the FeCO, as seen in other insects^47^, making it robust to transient manipulations of movement feedback alone. We also note that 9A neurons are embedded in complex VNC circuits beyond the circuit motif we identified. Therefore, manipulating 9A neurons may have off-target effects, making it inherently challenging to interpret causality experiments.^50^

### Behavioral function of sensory transmission in claw axons

We found that the position-encoding claw axons are not suppressed during active leg movements. Nonetheless, like hook axons, claw axons receive primarily GABAergic input and strongly express the GABA_A_ receptor gene Rdl. Most of the GABAergic input comes from interneurons belonging to the 19A hemilineage, which receive in part input from proprioceptive and tactile sensory neurons and in turn target primarily proprioceptive axons^36^. The function of this presynaptic inhibition remains to be determined. Based on studies in other animals, it could sharpen receptive fields through lateral inhibition, protect the sensory terminals from habituation, or reduce hysteresis in postsynaptic neurons^14^.

Overall, claw axons faithfully encoded position signals regardless of the behavioral context. During standing, the position feedback could contribute to postural reflexes^47^. During walking and grooming, the position feedback could help ensure proper leg placement in space^51–54^––a control task for which movement feedback is not suited. To support both motor functions, context-dependent modulation would have to occur at the level of interneurons^55^ rather than sensory axons. This is seen in the vestibular system of primates, where sensory axons faithfully relay head movement information regardless of the behavioral context, and modulation occurs at the level of interneurons^56^. A comprehensive analysis of the circuits downstream of claw and hook axons in the VNC connectomes would shed light on the specific pathways for leg motor control.

### Predictive inhibition from the brain

Our findings suggest that proprioceptive feedback inhibition is driven primarily by descending predictions from the brain. This conclusion is supported by: (1) 9A neurons being exclusively active during self-generated leg movements, (2) the chief 9A neuron receiving most of its input from descending neurons, some of which are known to drive walking and grooming, and (3) the strongest presynaptic descending neuron (web) releasing the chief 9A neuron from inhibition specifically during active leg movements. This is consistent with previous studies indicating that predictive inhibition originates in the brain. In primates, descending inputs drive the presynaptic inhibition of cutaneous and proprioceptive axons during active wrist movements^57,58^, although the specific source remains unknown. In zebrafish, an efference copy originating in the hindbrain inhibits the mechanosensory neurons of the lateral line^59,60^. In weakly electric fish, a cerebellum-like circuit in the brain cancels self-generated electrosensory input during swimming^61^. Notably, we found that the chief 9A neurons are targeted by several descending neurons, which in turn target many other neurons. This suggests that predictive inhibition of proprioceptive feedback results from multilayered descending input rather than dedicated prediction pathways.

Predictive inhibition can also originate in local motor circuits. The latter is seen in crickets, where the sensory axons of auditory neurons are suppressed during singing by the local motor circuits that produce the singing^62^. 9A neurons other than the chief 9A do receive input from premotor neurons in the VNC, so we cannot exclude a contribution from local motor circuits. However, these 9A neurons have far weaker connections with hook axons.

### Limitations of the study

The slow dynamics of the calcium signal relative to the high speed of fly leg movements prevented us from determining whether the activity of 9A or web neurons precedes movement. However, presynaptic inhibition effectively suppressed hook signals, which would require the presynaptic signal to be predictive in order to overcome sensorimotor delays. The calcium signals were also too slow to reveal whether modulation of sensory transmission is linked to the phase of the movement cycle. Phase-dependent modulation is common for spinal reflexes^3^, and also seen in insects. In walking locusts, proprioceptor axons receive a tonic inhibitory input that begins just prior to walking, and phasic inhibitory input during walking^16^. The proprioceptive axons of *Drosophila* are too small for intracellular electrophysiological recordings, but in the future, techniques like voltage imaging^63^ might enable recordings with sufficient temporal resolution to reveal the details of the feedback inhibition. In combination with mechanical perturbations of the leg, higher-resolution recordings could also reveal the extent to which externally-generated stimuli are transmitted to motor circuits, which would help clarify the function of the feedback inhibition.

## Conclusion

In this study, we found that proprioceptive movement feedback from the legs is selectively suppressed in behaving *Drosophila*. Selective context-dependent suppression is mediated by a segmentally-repeated circuit motif consisting of local GABAergic interneurons that are driven by descending neurons from the brain. In the future, it will be interesting to test whether the same logic extends to analogous movement-encoding (type Ia muscle spindle afferents) and position-encoding (type II muscle spindle afferents) proprioceptors in mammals.

## Acknowledgements

We thank members of the Tuthill laboratory for technical assistance and feedback on the manuscript, in particular Anne Sustar for confocal images and Su-Yee Lee for help with neural manipulation experiments. We thank Katharina Eichler and Gregory Jefferis for help with descending neuron identification. We also thank Jan M. Ache, Osama M. Ahmed, Eiman Azim, Eugenia M. Chiappe, and Lydia Zhang for feedback on the manuscript, Jasper S. Phelps, Wei-Chung Allen Lee, and the FANC community for their contributions to the proofreading of the VNC connectome, and James W. Truman for sharing *Drosophila* stocks. We used stocks obtained from the Bloomington *Drosophila* Stock Center (NIH P40OD018537). This work was supported by a Postdoctoral Research Fellowship from the Deutsche Forschungsgemeinschaft (DFG, German Research Foundation) project 432196121 and by the European Union’s Horizon Europe research and innovation program under the Marie Skłodowska-Curie grant agreement 101107596 to C.J.D, NIH grant K99NS117657 to S.A., as well as a Searle Scholar Award, a Klingenstein-Simons Fellowship, a Pew Biomedical Scholar Award, a McKnight Scholar Award, a Sloan Research Fellowship, the New York Stem Cell Foundation, and NIH grants R01NS102333 and U19NS104655 to J.C.T. J.C.T. is a New York Stem Cell Foundation – Robertson Investigator.

## Author contributions

C.J.D. and J.C.T. conceived of the study. C.J.D. developed the setup and analysis tools for calcium imaging in behaving animals. C.J.D. collected and analyzed calcium imaging data for sensory and 9A neurons in behaving animals. Y.L. collected and analyzed calcium imaging data for web neurons. S.A. collected and analyzed calcium imaging data for hook neurons during passive leg movements. G.M.C. collected and analyzed optogenetic data. C.J.D., S.A., and A.C. reconstructed neurons in FANC. C.J.D analyzed connectome and transcriptome data. C.J.D. visualized the results. C.J.D, S.A., and J.C.T. acquired funding. B.W.B. and J.C.T. supervised the project. C.J.D. and J.C.T. wrote the manuscript with input from other authors.

## STAR methods

### Key resources table

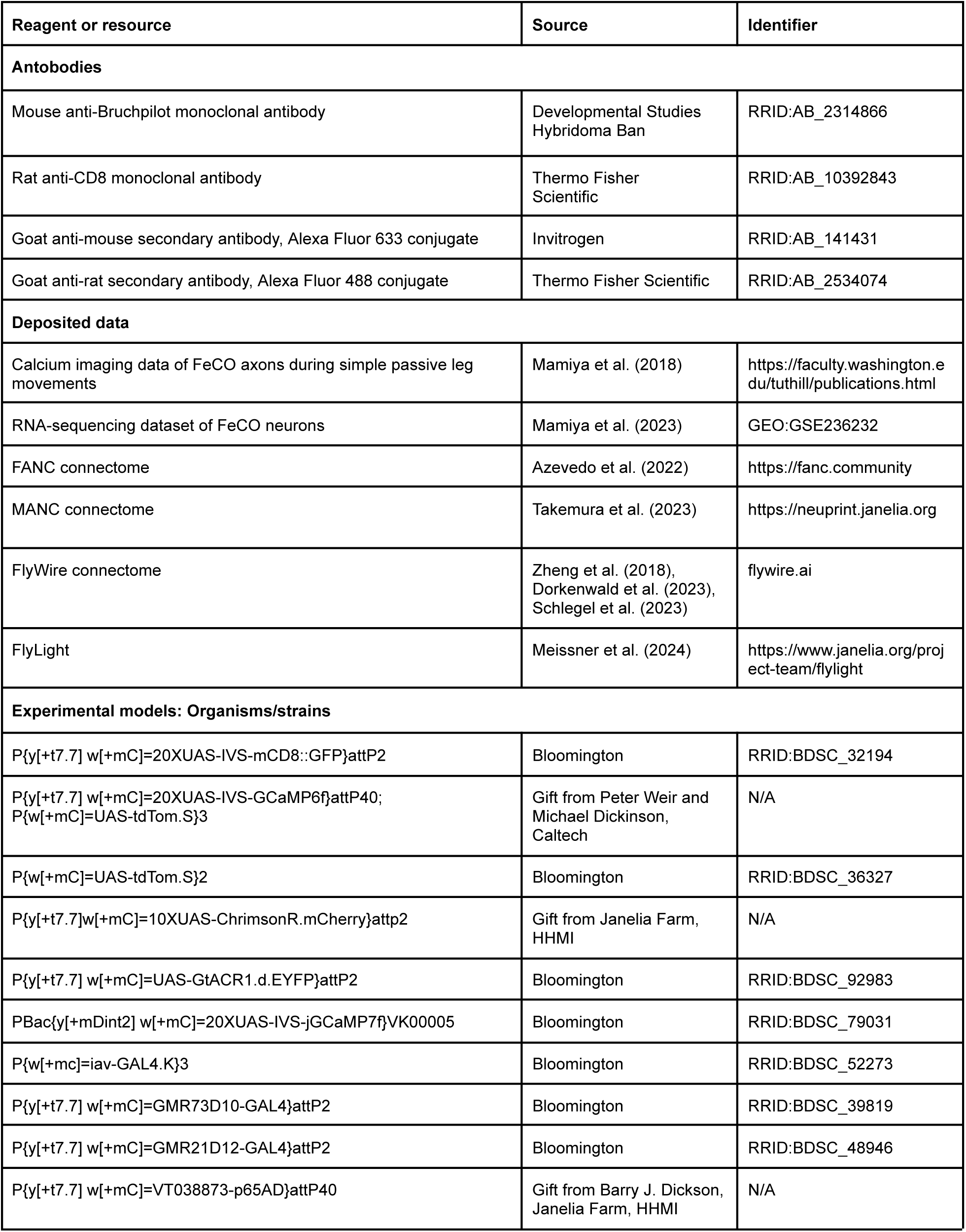
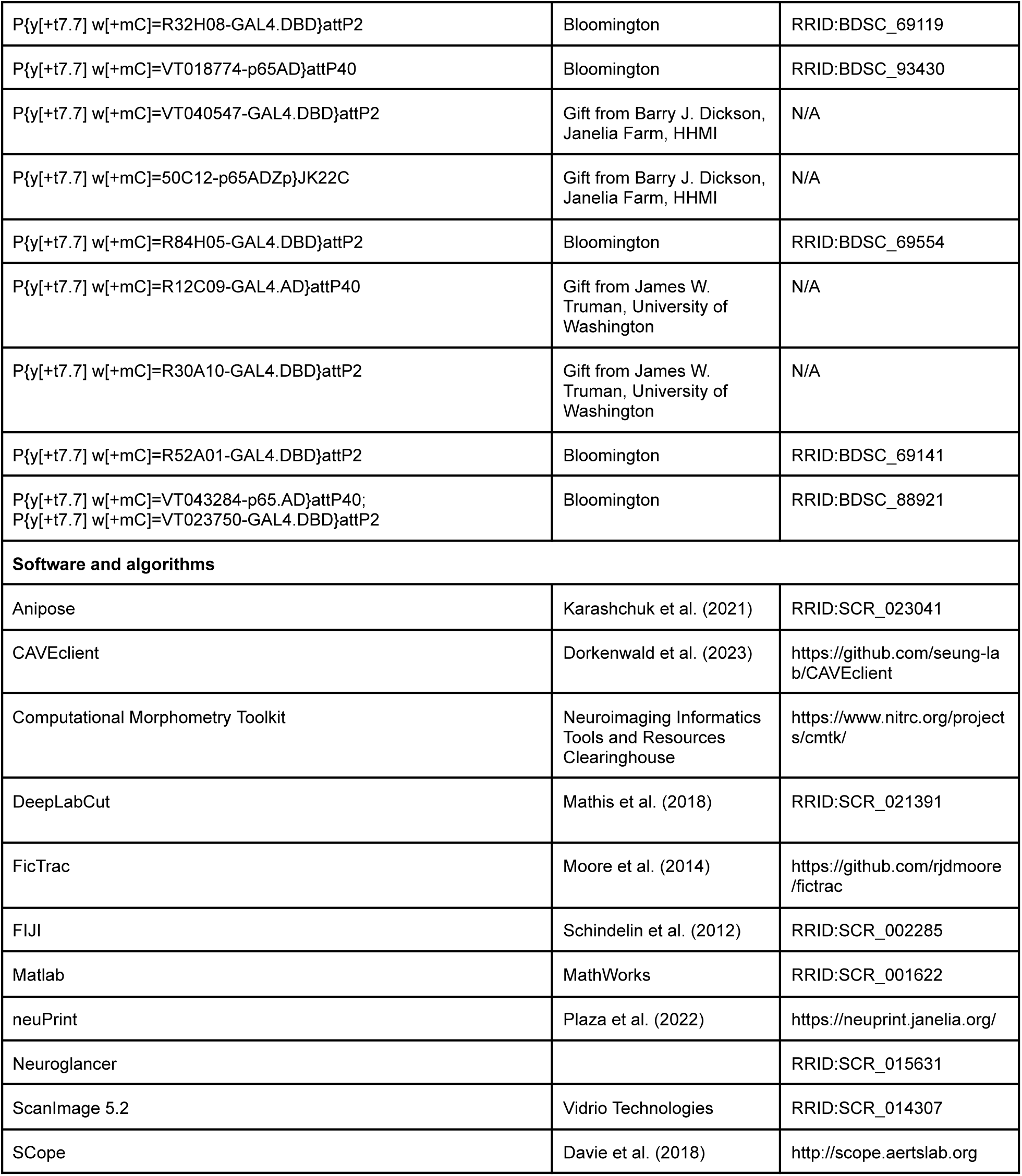

### Resource availability

#### Lead contact

Further information and requests for resources and reagents should be directed to and will be fulfilled by the lead contact, John C. Tuthill.

#### Materials availability

The split-GAL4 driver lines used in this study are available upon request from the lead contact. The underlying AD and DBD lines are listed in the key resources table and are available from the Bloomington *Drosophila* Stock Center or the lead contact.

#### Data and code availability

Calcium imaging and behavioral data generated for this paper will be available for download from Dryad. FANC connectome data were analyzed from the CAVE materialization version 840, timestamp 2024-01-17T08:10:01.179472. MANC connectome data were analyzed from version 1.0. FlyWire connectome data were analyzed from public release version 783. Analysis code used in this study will be available on GitHub (https://github.com/tuthill-lab). Any additional information required to reanalyze the data is available from the lead contact upon request.

### Experimental animals

We used *Drosophila melanogaster* raised on standard cornmeal and molasses medium at 25°C in a 14:10 hour light:dark cycle. We used flies 1 to 9 days post-eclosion. We used males for imaging experiments during behavior. We used females for the optogenetic experiments and the imaging experiments in the fully constrained preparation. For experiments involving optogenetic reagents, adult flies were placed on cornmeal agar with all-trans-retinal (35 mM in 95% EtOH; Santa Cruz Biotechnology) for 24-72 hours prior to the experiment. Vials were wrapped in foil to reduce optogenetic effects during development. The genetic driver lines used for each experiment are listed in a table below. In the neuromere of the left front leg, the claw line labels ∼25 axons, each hook line labels ∼5 axons, the club line labels ∼40 axons, the 9A line labels ∼10 neurons, and the web line labels 1 neuron. Claw, hook, and club neurons are cholinergic^18^. 9A neurons^28,36^ and the web neuron^36,44^ are GABAergic.

### Immunohistochemistry

For confocal imaging of FeCO axons and 9A neurons in the VNC, we crossed flies carrying the GAL4 driver to flies carrying pJFRC7-20XUAS-IVS-mCD8::GFP and dissected the VNC of females out of the thorax in *Drosophila* saline (103 mM NaCl, 3 mM KCl, 5 mM TES, 8 mM trehalose, 10 mM glucose, 26 mM NaHCO3, 1 mM NaH2PO4, 1.5 mM CaCl2, and 4 mM MgCl2; pH 7.1; osmolality adjusted to 270-275 mOsm). We fixed the VNC in a 4% paraformaldehyde PBS solution for 15 min. Next, we rinsed the VNC in PBS three times and put it in blocking solution (5% normal goat serum in PBS with 0.2% Triton-X) for 20 min, and then incubated it with a solution of primary antibody (rat anti-CD8 antibody, 1:50 concentration; mouse anti-Bruchpilot antibody for neuropil staining, 1:50 concentration) dissolved in blocking solution for 24 hours at room temperature. At the end of the first incubation, we washed the VNC with PBS with 0.2% Triton-X (PBST) three times, and then incubated the VNC in a solution of secondary antibody (goat anti-rat antibody Alexa Fluor 488, 1:250 concentration; goat anti-mouse antibody Alexa Fluor 633, 1:250 concentration) dissolved in blocking solution for 24 hours at room temperature. Finally, we washed the VNC in PBST three times and then mounted it on a slide with Vectashield (Vector Laboratories). We acquired z-stacks of each VNC on a confocal microscope (Zeiss 510; Zeiss). We aligned the expression pattern in the VNC using the Computational Morphometry Toolkit (CMTK; Neuroimaging Informatics Tools and Resources Clearinghouse) to a female VNC template^64^ in Fiji^65^. Confocal stacks of the claw line, one of the hook flexion lines (driver line 1), and the web line were downloaded from the GAL4 and split-GAL4 collections on FlyLight^66,67^.

### Analysis of RNA-sequencing data

The RNA-sequencing data of FeCO neurons was generated in a previous study^19^. We queried the dataset for expression of different receptor genes in claw and hook neurons using SCope^68^.

### Fly preparation for *in vivo* two-photon calcium imaging in the VNC

For recording calcium signals in the VNC of behaving flies, we adapted a previously described preparation^69,70^. We clipped the fly’s wings under cold anesthesia. Then, we pushed the dorsal part of the thorax through a hole (0.8 mm width, 0.95 mm length) in a curved steel sheet at the bottom side of a custom-made holder. The thorax was fixed using UV-curing glue (KOA 300; Kemxert) applied around the perimeter of the thorax on the top side of the holder. This left the fly’s legs, head, and abdomen on the bottom side of the holder free to move. The top side of the holder was then immersed in *Drosophila* saline. To gain optical access to the VNC, a rectangular piece of cuticle was removed from the dorsal thorax. This exposed the indirect flight muscles (IFMs) while leaving the body-wall muscles intact. IFMs were parted along the midline of the body using a tapered insect pin (0.1 mm diameter; Living Systems Instrumentation). We waited ∼60 min for IFMs to partly dissolve. Remaining IFMs were then removed using the insect pin. Removing IFMs exposed the proventriculus (or cardia, a part of the gut) and surrounding tracheae above the neuromeres of the front legs. Fine forceps were used to pull out the anterior trachea above the neck connective and remove the underlying fat bodies. Then, the proventriculus was moved to the right side of the thorax using an insect pin, exposing most of the neuromere of the left front leg. The pin was held in place by a sculpting compound (Super Sculpey Firm) positioned next to the thorax. A second insect pin was inserted into the sculpting compound and thorax to displace the left lateral trachea. Care was taken not to touch the VNC while displacing the gut and tracheae. Leaving the gut and tracheae intact proved critical for normal fly behavior and allowed us to record from the VNC for several hours. After the dissection, the fly holder was mounted onto a three-axis manipulator, and the fly was positioned above the treadmill. We typically gave flies 30-60 min to recover from the preparation before starting the experiments.

For recording calcium signals in the VNC while controlling tibia position with the magnet-motor system (see below), we used a previously described preparation in which the fly is oriented ventral side up^18^. We first cold anesthetized the fly on ice and then pushed the head and ventral thorax through a hole in a steel sheet of a custom-made holder. The head and thorax were fixed using UV-curing glue (KOA 300; Kemxert). The abdomen and legs were placed on the bottom side of the holder. To control the femur-tibia joint angle, we glued the femur of the right front leg to the bottom side of the holder and attached a small piece of insect pin (∼1 mm length, 0.1 mm diameter; Living Systems Instrumentation) to the tibia and tarsus. The pin was painted black (Super Black; Speedball Art Products) to improve image contrast for movement tracking (see below). All other legs were glued to the holder to not interfere with the movement of the tibia of the right front leg. The top side of the holder was then immersed in *Drosophila* saline. To gain optical access to the VNC, the cuticle covering the front leg neuromeres was removed with fine forceps. Fat bodies and larger trachea covering the imaging region of interest were removed as well. Finally, we removed the digestive tract with fine forceps to reduce the movement of the VNC.

### *In vivo* two-photon image acquisition

For recording calcium signals in the VNC during behavior and motor-controlled movements of the tibia, we used two two-photon Movable Objective Microscopes (MOM; Sutter Instruments) with a 20x water-immersion objective (Olympus XLUMPlanFI, 0.95 NA, 2.0 mm wd; Olympus) and a 40x water-immersion objective (0.8 NA, 2.0 mm wd; Nikon Instruments), respectively. Neurons of interest expressed the calcium indicator GCaMP6f or GCaMP7f (green fluorescence) and the structural marker tdTomato (red fluorescence). Fluorophores were excited at 920 nm by a mode-locked Ti:sapphire laser (Chameleon Vision S; Coherent). We used a Pockels cell to keep the power at the back aperture of the objective below ∼35 mW. Emitted fluorescence was directed to two high-sensitivity GaAsP photomultiplier tubes (Hamamatsu Photonics) through a 705 nm edge dichroic beamsplitter followed by a 580 nm edge image-splitting dichroic beamsplitter (Semrock). Fluorescence was band-passed filtered by either a 525/50 (green) or 641/75 (red) emission filter (Semrock). Image acquisition was controlled with ScanImage 5.2 (Vidrio Technologies) in Matlab (MathWorks). Each microscope was equipped with a galvo-resonant scanner, and each objective was mounted onto a piezo actuator (Physik Instrumente; digital piezo controller E-709). For recordings during behavior, we acquired volumes of three 512 x 512 pixel images spaced 5 μm apart in depth (10 μm total) at a speed of 8.26 volumes per second. We typically recorded 400 volumes (∼50 s) per trial. For recordings during motor-controlled movements of the tibia, we acquired volumes of three 256 x 128 pixel images spaced 10 µm apart in depth (20 µm total) at a speed of 36.7 volumes per second. Previous experiments revealed that calcium signals in claw and hook axons do not differ qualitatively across different axon branches when the leg is passively moved^18^. Therefore, we focused our experiments on a single imaging region. All experiments were performed in the dark at room temperature.

### Two-photon calcium imaging analysis

Two-photon images were analyzed with custom scripts in Matlab. Images acquired during behavior were analyzed in nine steps. First, we smoothed each image with a Gaussian filter (sigma = 3 pixels; size = 5 x 5 pixels). Second, we corrected for horizontal movement of the VNC. Each tdTomato image was aligned to the average tdTomato signal of the recorded trial via translations using a cross-correlation-based image registration algorithm^71^ (upsampling factor = 4). The same translations were then applied to the GCaMP images. Third, the three GCaMP and tdTomato images per volume were averaged. Fourth, we extracted the mean fluorescence in manually drawn regions of interest (ROIs). Fifth, we corrected for vertical movement of the VNC by computing the ratio of GCaMP fluorescence to tdTomato fluorescence in each frame^72^. Dividing the GCaMP fluorescence by the tdTomato fluorescence decreased the impact of vertical movement, because such movements result in correlated changes in both signals, and the tdTomato signal is independent of neural activity. Sixth, to facilitate comparisons across trials and flies, ratio values were z-scored by subtracting the mean of a baseline ratio and dividing by the standard deviation of that baseline ratio. The baseline was defined in each trial as the 10% smallest ratio values. Seventh, z-scored ratio values were upsampled to the sampling rate of leg tracking (300 Hz) using cubic spline interpolation. Eighth, upsampled ratio values were low-pass filtered using a moving average filter with a time window of 0.2 s. Ninth, to facilitate comparisons with predicted calcium values, we normalized the measured calcium values to be between zero and one by dividing each z-scored ratio value by the maximum z-scored ratio value for a given genetic driver line in the dataset.

Two-photon images acquired during motor-controlled movements of the tibia were analyzed similarly, but due to a lack of VNC movement in that setup, correcting for horizontal and vertical movement of the VNC (steps 2 and 5 above) was not necessary. Instead, we computed the change in GCaMP fluorescence relative to a baseline per trial. For each frame, we subtracted the mean of the baseline from the GCaMP fluorescence and divided by the mean of the baseline. The baseline was defined per trial as the lowest average GCaMP fluorescence in a window of 0.27 s (10 frames). Then, calcium signals were z-scored, upsampled, low-pass filtered, and normalized as described above (steps 6–9).

To fit the computational models (see below), we z-scored and normalized the calcium imaging data from Mamiya et al.^18^ in the same manner as the data recorded from behaving flies. That is, we first z-scored the calcium signals per trial relative to a baseline, and then divided each z-scored value by the maximum z-scored value for a given genetic driver line in the dataset.

### Treadmill for calcium imaging experiments

The omnidirectional treadmill consisted of a patterned Styrofoam ball (9.1 mm diameter; 0.12 g) floating on air in an aluminum holder. The air flow was set to ∼500 ml/min. The ball was illuminated by two infrared LEDs (850-nm peak wavelength; ThorLabs) via optical fibers. Ball movements were recorded at 30 Hz with a camera (Basler acA1300-200um; Basler AG) equipped with a macro zoom lens (Computar MLM3X-MP; Edmund Optics). Ball rotations around the fly’s cardinal body axes (forward, rotational, sideward) were reconstructed offline using FicTrac^73^. Rotational velocities of the fly were calculated based on the known diameter of the ball. Velocities were upsampled to the sampling rate of leg tracking (300 Hz) using cubic spline interpolation and low-pass filtered using a moving average filter with a time window of 0.2 s. The treadmill was mounted onto a one-axis manipulator. This allowed us to remove the treadmill in between trials and record data for leg movements in the air or on the platform.

### Moveable platform for calcium imaging experiments

The platform consisted of a metal pin (0.5 mm diameter, 4.4 mm length) mounted onto a three-axis micromanipulator (MP-225; Sutter Instruments). The pin was wrapped in black sandpaper to provide sufficient grip for the flies’ tarsi. The micromanipulator was controlled manually.

### Magnet-motor system for moving the tibia

We used a previously described magnet-motor system^18^ to control the femur-tibia angle during calcium imaging. We moved the tibia/pin to different positions via a cylindrical rare earth magnet (10 mm height, 5 mm diameter). The magnet was attached to a steel post whose position was controlled with a programmable servo motor (SilverMax QCI-X23C-1; QuickSilver Controls). The motor was mounted onto a micromanipulator (MP-225; Sutter Instruments). This allowed us to adjust the motor position so that the magnet moved in a circular trajectory centered at the femur-tibia joint.

The movement of the magnet, and with that, the tibia, was controlled with a custom script in Matlab. We imposed movements that were representative of femur-tibia joint angles and velocities recorded during walking and grooming (Figure S5B). Each trial started at a femur-tibia angle of ∼90°. In “walking”’ trials, we replayed 67 movement bouts containing different front leg walking kinematics. Each bout was 2 s in duration. In “grooming”’ trials, we replayed 12 movement bouts containing different front leg grooming kinematics. Bout durations ranged from 0.3 s to 2 s. Because movement bouts did not necessarily start or end at a femur-tibia angle of 90°, we added a 0.25 s transition phase before and after each bout, in which the tibia was linearly moved to the start position or back to 90°. The tibia was not moved for 0.5 s in between stimuli.

### Tracking of the femur-tibia joint during calcium imaging experiments

For recordings during behavior, movements of the left front leg were recorded at 300 Hz with two cameras (Basler acA800-510um; Basler AG) equipped with 1.0x InfiniStix lenses (68 mm wd; Infinity) and 875 nm short pass filters (Edmund Optics). The leg was illuminated by an infrared LED (850-nm peak wavelength; ThorLabs) via an optical fiber. We trained a deep neural network (DeepLabCut^74^) to automatically track all leg joints in each camera view. 2D tracking data from both camera views were then combined to reconstruct leg joint positions and angles in 3D using Anipose^75^. Specifically, we applied a median filter on the 2D tracking data and then used spatiotemporally regularized triangulation. The two cameras were calibrated using a ChArUco board (6x6 markers, 4 bits per marker, 0.125 mm marker length). 3D leg tracking was necessary to provide accurate femur-tibia joint angle information for the computational models that predicted calcium signals in our neurons of interest (see below).

For recordings with the magnet-motor system, movements of the right front leg tibia were recorded at 200 Hz with a single camera (Basler acA800-510um; Basler AG) equipped with a 1.0x InfiniStix lens (94 mm wd; Infinity) and a 900 nm short pass filter (Edmund Optics). Because the servo motor was placed directly under the fly, we placed the camera to the side and used a prism (Edmund Optics) to capture the view from below. The leg was illuminated by an infrared LED (850-nm peak wavelength; ThorLabs) via an optical fiber. The coxa-femur, femur-tibia, and tibia-tarsus joints were tracked using DeepLabCut. Because the tibia moved in a single plane parallel to the surface of the holder, 2D tracking was sufficient to provide accurate femur-tibia joint angle information for the computational models.

### Data annotations and data selection for calcium imaging experiments

For trials involving the treadmill and platform, fly behavior was classified semi-automatically based on the leg tracking data. All classifications were reviewed and manually corrected if necessary. First, movement of the left front leg was determined based on the speed of the leg’s tarsus in the side-view camera. The velocity was low-pass filtered using a moving average filter with a time window of 0.3 s. Frames in which the velocity exceeded 0.8 px/s were classified as moving. The resulting binary behavioral sequence was low-pass filtered by removing epochs shorter than 250 ms (hysteresis filter). That is, the behavioral sequence could only change state if at least 250 ms were in a new state. In trials involving the treadmill, movements were further classified as walking or grooming based on the movement of the left middle leg. Movement of the middle leg was determined analogously to that of the front leg (low-pass filter followed by thresholding and hysteresis filter). Epochs in which both the front leg and the middle leg moved were classified as walking. Epochs in which the front leg but not the middle leg moved were classified as front leg grooming. Front leg movements other than walking or grooming (e.g., extended downward pushing) were manually classified as “other.”

For trials involving the platform, we additionally manually annotated periods of passive leg movement based on the leg videos. For hook flexion neurons, we annotated passive flexions of the femur-tibia joint (Figure 3G). For hook extension neurons, we annotated passive extensions of the femur-tibia joint (Figure S4H). For 9A neurons, we annotated all passive movements of the leg (Figure 4H). For web neurons, we manually annotated hind leg grooming events for the computational model (see below).

Frames were manually excluded from the analysis if the front leg was involved in movements other than walking or grooming on the treadmill (e.g., extended downward pushing), the femur-tibia joint of the front leg was not tracked correctly, or the two-photon image registration failed (e.g., the VNC moved out of the imaging volume). When calculating the cross-correlation for hook axon recordings during passive leg movements with the platform, we additionally excluded frames in which the leg moved actively. Behavioral bouts and movement transitions were excluded if they were shorter than the desired minimum duration (see figure legends).

For some genotypes (see table of genotypes), we recorded neural activity with GCaMP6f and GCaMP7f. We did not observe any differences in the calcium signals and therefore pooled recordings for our analysis.

### Computational models for predicting calcium signals in neurons

We constructed computational models to predict time courses of calcium signals in claw, hook, club, 9A, and web neurons from time courses of femur-tibia joint kinematics or binary behavior variables. The time courses were fed into a neuron type-specific activation function, which was convolved with a double exponential function to mimic the temporal dynamics of GCaMP:

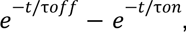

with an onset time constant τ_on_ = 0.03 s and an offset time constant τ_off_ = 0.3 s. The time constants were tuned to match the measured calcium signals in claw and hook axons in Mamiya et al.^18^ (Figure S2).

The activation functions for claw and hook neurons were based on calcium imaging and leg tracking data from Mamiya et al.^18^, where the femur-tibia joint was passively moved using ramp-and-hold stimuli (Figure S2). In that dataset, calcium signals in claw axons are lowest at a joint angle of 90° and increase non-linearly with increasing flexion or extension. To model this encoding, we first subtracted 90° from the tracked femur-tibia joint angle. Then, we fitted a 4th order polynomial activation function (convolved with the GCaMP kernel) to the z-scored and normalized calcium signals using nonlinear least-squares optimization (lsqcurvefit; Matlab; Figure S2A). In our dataset, calcium signals were weakest at a joint angle of 80° (Figure 2F). Thus, for our dataset, we subtracted 80° from the tracked femur-tibia joint angle, but used the same activation function. The 10° difference between the datasets is likely related to differences in leg tracking, not encoding.

Hook neurons were assumed to encode flexion or extension direction. To model this encoding, the joint angle velocity was fed into a binary step function (Figure S2A). For hook flexion neurons, we used a threshold of −5 deg/s for the dataset from Mamiya et al.^18^ and −50 deg/s for our dataset. Different thresholds were chosen to account for different amounts of tracking noise in the datasets. Mamiya et al.^18^ did not test hook extension neurons. Based on a recent study^19^, hook extension neurons have the opposite encoding of hook flexion neurons, which we modeled with a binary step function with a threshold of 50 deg/s in our dataset (Figure S2A).

Club neurons were assumed to encode bidirectional movement^18^. To model this encoding, the joint angle velocity was fed into a rectangular function with thresholds of ±50 deg/s (Figure S2A).

9A neurons were assumed to encode bidirectional movement as well, which we modeled with the same activation function that we used for club neurons (Figure S2A). However, we assumed that 9A neurons do not respond to passive leg movements. To model this, the joint angle velocity input was set to zero during passive leg movements.

To model the activity of the web neuron, we used a binary vector indicating when all legs were at rest as activation function. This required manual annotations for legs other than the left front leg (see above).

To facilitate comparisons with measured calcium values, we normalized the predicted calcium values to be between zero and one by subtracting the minimum predicted value in each trial and dividing by the maximum predicted value for a given genetic driver line in the dataset.

### Fly preparation for optogenetic experiments

We clipped the flies’ wings under cold anesthesia just prior to experiments in order to increase walking and prevent visual obstruction of the legs and thorax. To position the fly above the spherical treadmill, a tungsten wire was attached to the dorsal thorax with UV-curing glue (KOA 300; Kemxert).

### Setup for optogenetic stimulation

We coaxed flies to walk on a treadmill by displaying visual stimuli on a semi-circular green LED display^20,32^. We displayed a single dark bar (30° width) on a light background, and sinusoidally oscillated the bar at 2.7 Hz across 48.75° about the center of the fly’s visual field. During periods between trials, the LED panels displayed a fixed dark stripe (30°) on a bright background in front of the tethered fly. To optogenetically activate or silence 9A neurons, we used a red laser (638 nm, Laserland) or green laser (532 nm, CST DPSS laser, Besram Technology), respectively. The lasers were pulsed at 1200 Hz with a 60% duty cycle and a resulting power of ∼80 mW/mm^2^ at the target. The lasers were aimed at the ventral thorax at the base of the left front leg^20,32^. Previous experiments indicate that this optogenetic stimulation primarily affects neurons in the neuromere of the left front leg^32^, though we cannot rule out effects on other VNC neurons. Fly behavior was recorded in 2 s trials. The laser stimulus began at 0.5 s and lasted 1 s.

### Treadmill for optogenetic experiments

The treadmill was the same as for the calcium imaging experiments. Ball movements were recorded at 30 Hz with a camera (FMVU-03MTM-CS; Point Grey Research) equipped with a macro zoom lens (Computar MLM3X-MP; Edmund Optics). Ball rotations around the fly’s cardinal body axes (forward, rotational, sideward) were reconstructed from live video using FicTrac^73^.

### Tracking of leg joints during optogenetic experiments

We used a previously described camera setup^75^ to record the movements of all legs during optogenetic experiments. Six cameras (Basler acA800-510; Basler AG) equipped with a macro zoom lens (Computar MLM3X-MP; Edmund Optics) were evenly distributed around the fly, providing full video coverage of all six legs. We used a previously trained DeepLabCut network^75^ to automatically track all leg joints in each camera view. 2D tracking data from all camera views were then combined to reconstruct leg joint positions and angles in 3D using Anipose^75^. Step cycles were classified automatically based on thresholds on the velocity of the leg tips as described previously^34^.

### Reconstruction of FeCO axons and presynaptic neurons in the FANC connectome

Neurons in the Female Adult Nerve Cord (FANC) electron microscopy dataset^76^ were previously segmented in an automated manner^23^. To manually correct the automated segmentation of claw and hook axons and their presynaptic neurons, we used Google’s collaborative Neuroglancer interface (https://github.com/google/neuroglancer). Pre- and postsynaptic neurons that made less than three synapses with a neuron of interest were excluded from connectivity analyses. Neuron annotations were managed by the Connectome Annotation Versioning Engine (CAVE)^77^. We used custom scripts in Python to interact with CAVE via CAVEclient^77^ and analyze connectivity.

### Identification of hemilineages and fast-acting neurotransmitters in the FANC connectome

In *Drosophila*, neurons that share a developmental origin (i.e., belong to the same hemilineage) possess common anatomical features^27^ and release the same fast-acting neurotransmitter^28^ (GABA, glutamate, or acetylcholine). We took advantage of this knowledge to identify the hemilineage and thus the fast-acting neurotransmitter of each VNC neuron presynaptic to the claw and hook axons in the FANC connectome. For identification, we relied on light microscopy images of sparse GAL4 lines^28,78^, cell body position along the dorsal-ventral axis, and personal communication (James W. Truman, David Shepherd, Haluk Lacin, and Elizabeth Marin).

### Definition of neuron types in the FANC connectome

Neurons presynaptic to the 9A neurons in the FANC connectome were identified as sensory neurons, descending neurons, or premotor neurons. Sensory neurons had processes entering the VNC from peripheral nerves and no cell body in the VNC. Descending neurons had a process in the neck connective and no cell body in the VNC. Premotor neurons were presynaptic to leg motor neurons in the neuromere of the left front leg. These neurons were previously annotated by Lesser et al.^79^.

### Circuit analysis in the MANC connectome

The Male Adult Nerve Cord (MANC) connectome^35^ and its annotations^36,40^ were queried via the neuPrint API and web explorer^80^. Neurons that made less than five synapses with a neuron of interest were excluded from connectivity analyses.

### Circuit analysis in the FlyWire connectome

The FlyWire connectome^42,43^ and its annotations^44^ were queried with custom Python scripts via CAVEclient^77^. Neurons that made less than five synapses with the web neuron were excluded from connectivity analyses. To analyze the inputs that the web neuron receives from different brain neuropils, we first calculated the relative number of input synapses of each presynaptic neuron that are located in the different brain neuropils (pooled across hemispheres). These relative synapse counts per presynaptic neuron were then weighted by multiplication with the number of synapses between the presynaptic neuron and the web neuron. Finally, we summed the weighted synapse counts per neuropil and expressed them relative to the sum of weighted synapse counts across neuropils.

### Table of genotypes

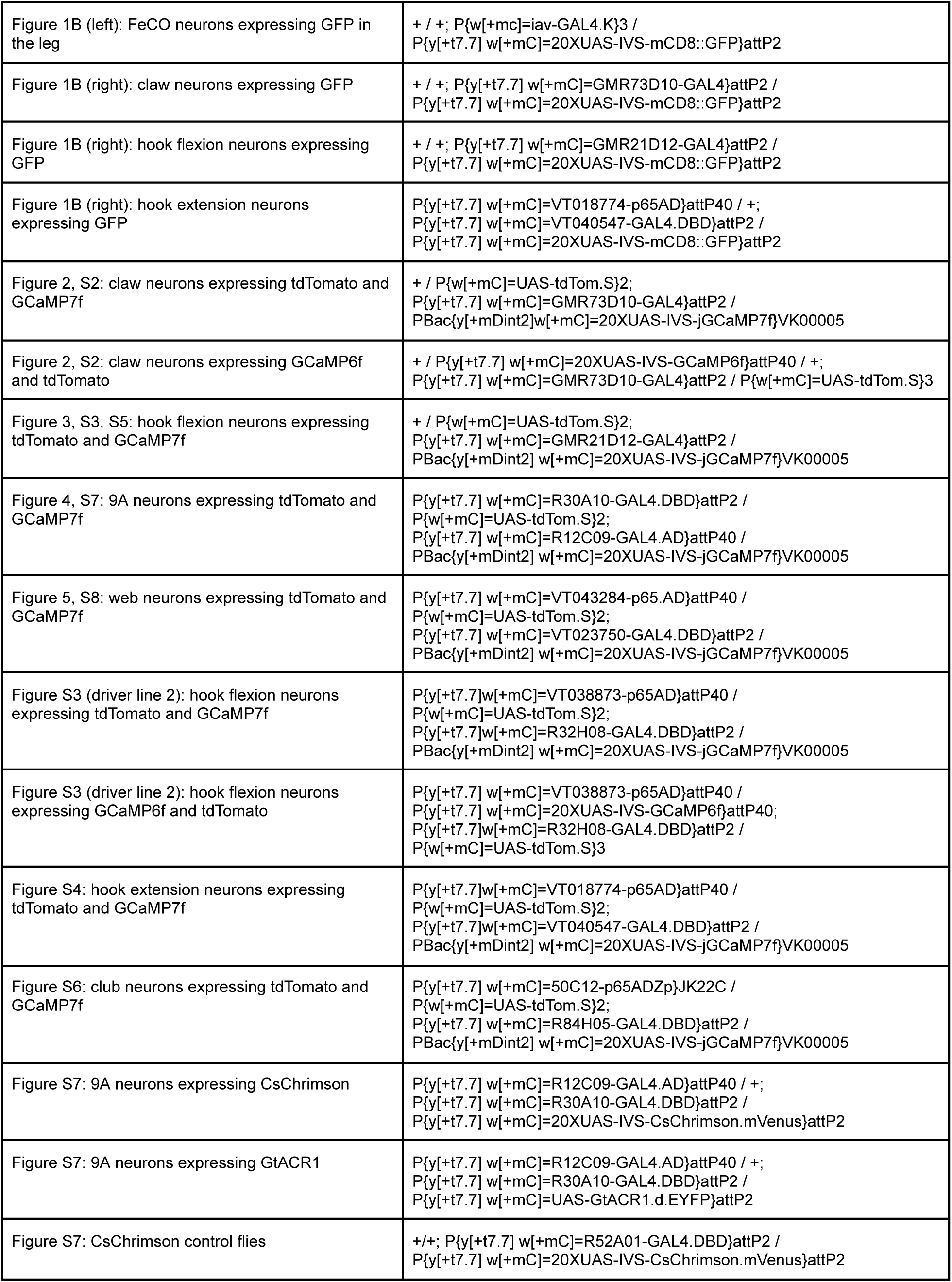
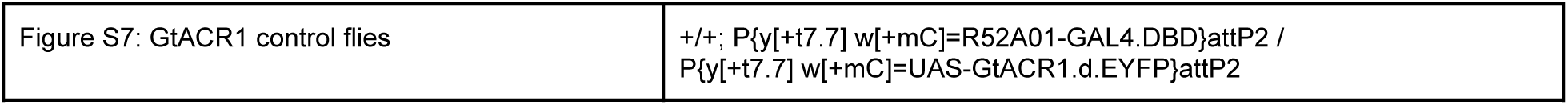

