## Supplementary Figures and Tables for "Presynaptic inhibition selectively suppresses leg proprioception in behaving *Drosophila*"

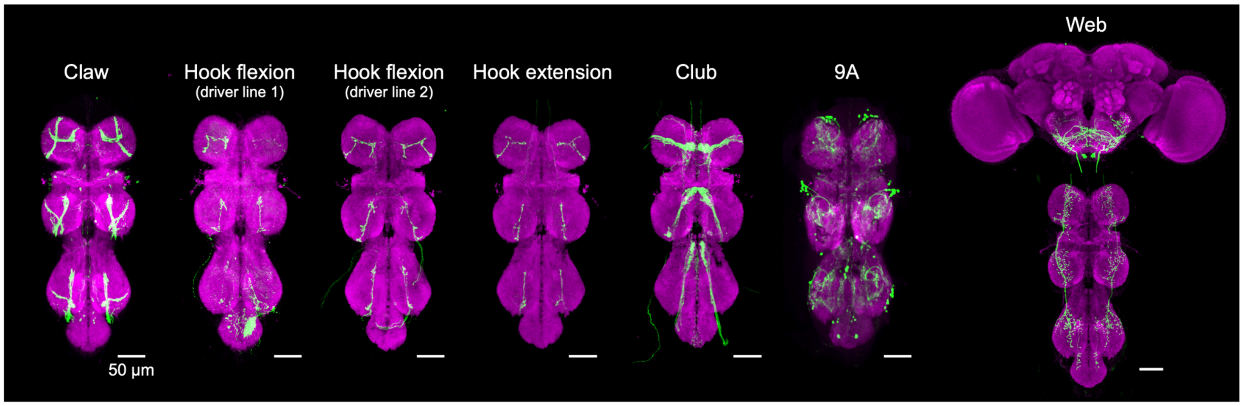


**Figure S1. Related to Figures 2-5**
Confocal images showing the expression patterns of the GAL4 and split-GAL4 driver lines used to label FeCO neurons, 9A neurons, and web neurons. Green: GFP or mVenus; magenta: neuropil stain (nc82). Genotypes are listed in the table of genotypes. Images of the claw line, the hook flexion line 1, and the web line are from FlyLight.

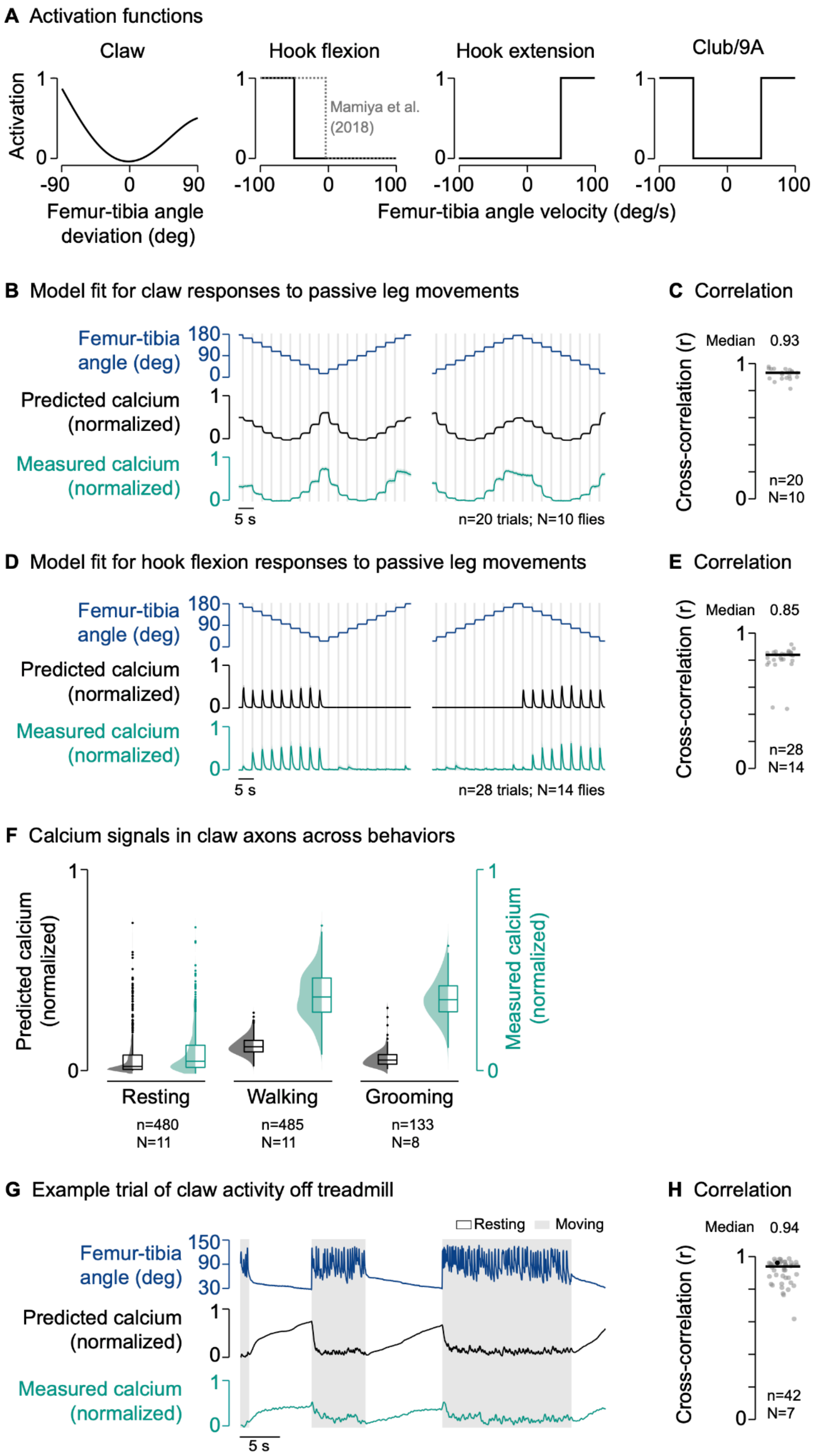

A Activation functions
Claw
Hook flexion
Hook extension
Club/9A
Activation
Mamiya et al. (2018)
Femur-tibia angle deviation (deg)
Femur-tibia angle velocity (deg/s)
B Model fit for claw responses to passive leg movements
Femur-tibia angle (deg)
Predicted calcium (normalized)
Measured calcium (normalized)
5 s
n=20 trials; N=10 flies
C Correlation
Cross-correlation (r)
Median 0.93
n=20
N=10
D Model fit for hook flexion responses to passive leg movements
Femur-tibia angle (deg)
Predicted calcium (normalized)
Measured calcium (normalized)
5 s
n=28 trials; N=14 flies
E Correlation
Cross-correlation (r)
Median 0.85
n=28
N=14
F Calcium signals in claw axons across behaviors
Predicted calcium (normalized)
Measured calcium (normalized)
Resting
n=480
N=11
Walking
n=485
N=11
Grooming
n=133
N=8
G Example trial of claw activity off treadmill
Resting
Moving
Femur-tibia angle (deg)
Predicted calcium (normalized)
Measured calcium (normalized)
5 s
H Correlation
Cross-correlation (r)
Median 0.94
n=42
N=7

**Figure S2. Related to Figure 2**

(A) Activation functions for claw, hook flexion, hook extension, and club and 9A neurons.

(B) Measured and predicted (fitted) calcium signals of claw axons in response to applied ramp-and-hold movements of the femur-tibia joint. Experimental data from Mamiya et al. (2018). Lines show mean of animal means, shadings show standard error of the mean. n: number of trials (10 trials per ramp-and-hold stimulus, totalling 20 trials for both stimuli); N: number of flies.

(C) Cross-correlation coefficient between predicted and measured calcium signals per trial at a time lag of zero. The black line shows the median. n: number of trials; N: number of flies.

(D) Same as (B) but for hook flexion axons.

(E) Same as (C) but for hook flexion axons.

(F) Median predicted and measured calcium signals in claw axons during resting, walking, and grooming. Bouts are ≥1 s in duration. Distributions show kernel density estimations. n: number of behavioral bouts; N: number of flies. Note that the predicted calcium signals during walking and grooming appear weak because they are normalized to the maximum predicted value in the dataset, which occurred during an outlier resting bout in which the leg was extremely flexed.

(G) Example trial of two-photon calcium imaging of claw axons in the neuromere of the left front leg and behavior tracking without the treadmill.

(H) Same as (C) but for claw axons imaged without the treadmill. The black dot marks the trial shown in (G).

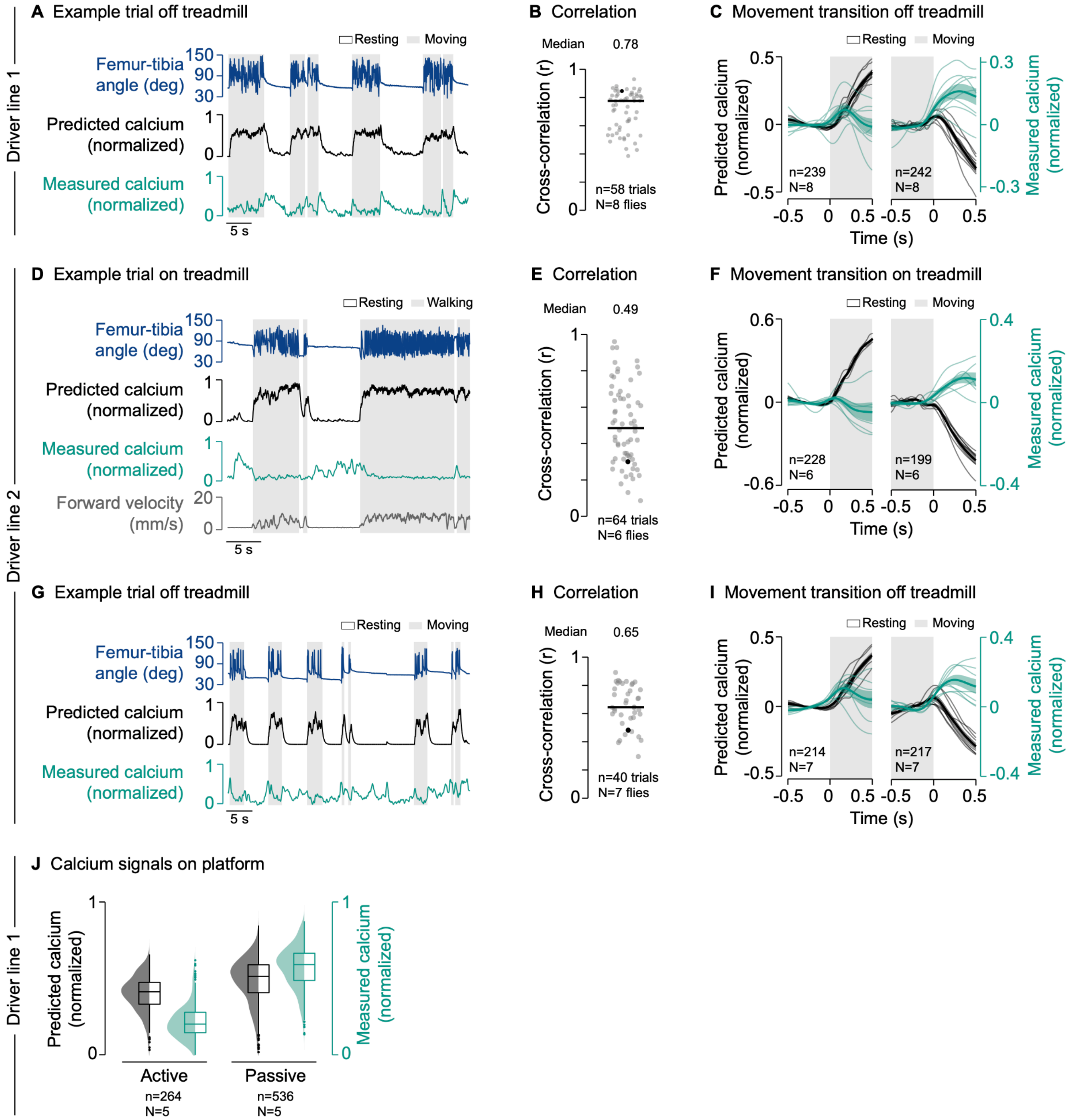


**Figure S3. Related to Figure 3**

(A) Example trial of two-photon calcium imaging of hook flexion axons in the neuromere of the left front leg and behavior tracking without the treadmill.

(B) Cross-correlation coefficient between predicted and measured calcium signals per trial at a time lag of zero. The black line shows the median. The black dot marks the trial shown in (A). n: number of trials; N: number of flies.

(C) Predicted and measured calcium signals aligned to the transitions into and out of movement. Signals are baseline subtracted (mean from -0.5 to 0 s). Thin lines show animal means, thick lines show mean of means, shadings show standard error of the mean. n: number of transitions; N: number of flies.

(D) Example trial of two-photon calcium imaging of hook flexion axons (second driver line) in the neuromere of the left front leg and behavior tracking on the treadmill.

(E) Same as (B) but for hook flexion axons (second driver line) imaged on the treadmill.

(F) Same as (C) but for hook flexion axons (second driver line) imaged on the treadmill. Movement includes walking and grooming.

(G) Example trial of two-photon calcium imaging of hook flexion axons (second driver line) in the neuromere of the left front leg and behavior tracking without the treadmill.

(H) Same as (B) but for hook flexion axons (second driver line) imaged without the treadmill.

(I) Same as (C) but for hook flexion axons (second driver line) imaged without the treadmill.

(J) Median predicted and measured calcium signals during active and passive movement bouts on the platform. Bouts are ≥0.5 s in duration. Distributions show kernel density estimations. n: number of movement bouts; N: number of flies.

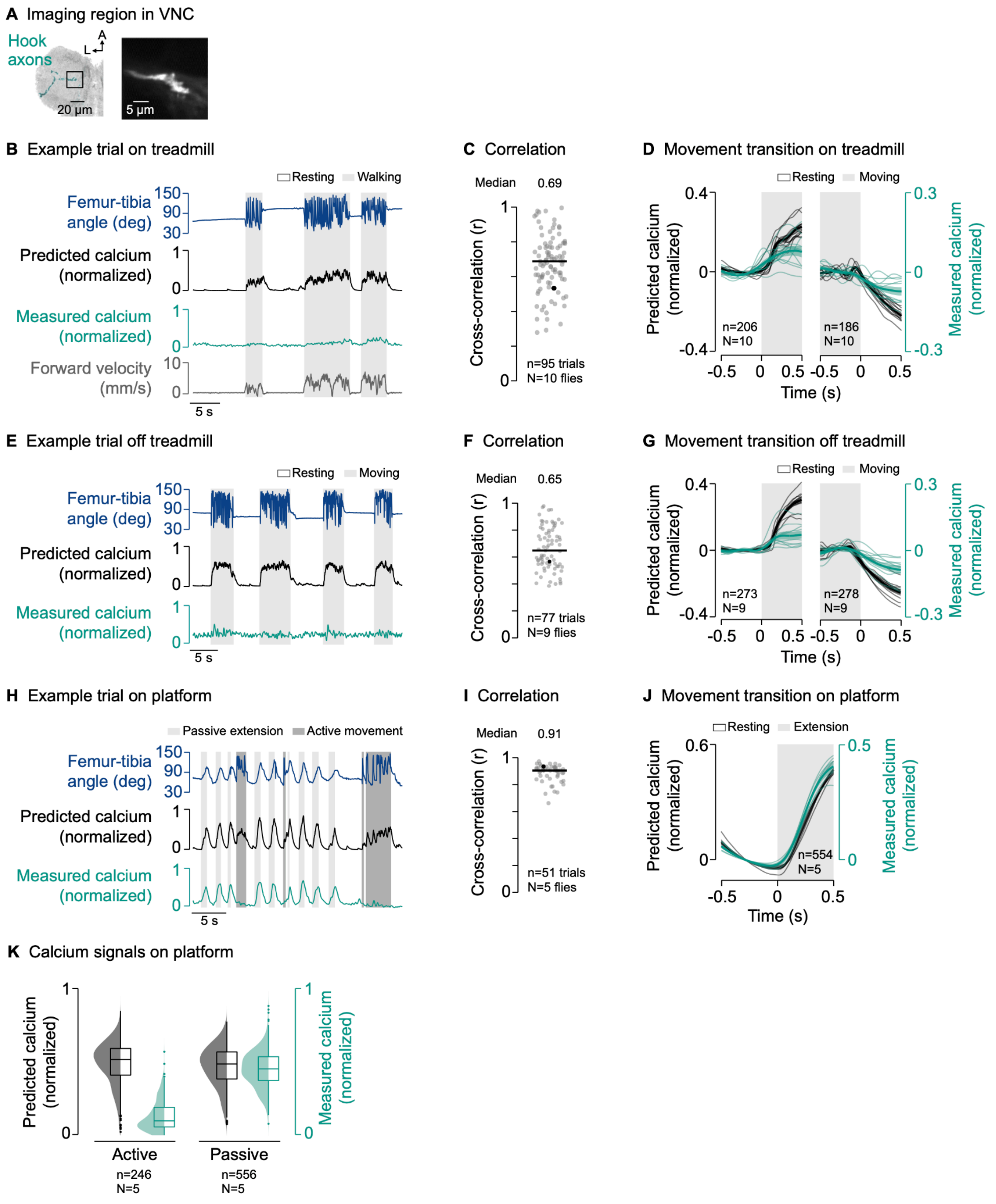


**Figure S4. Related to Figure 3**

(A) Left: Confocal image of hook extension axons in the neuromere of the left front leg. The black box indicates the imaging region. Green: GFP; gray: neuropil stain (nc82). A: anterior; L: lateral. Right: Mean tdTomato signal within the imaging region during an example trial.

(B) Example trial of two-photon calcium imaging of hook extension axons in the neuromere of the left front leg and behavior tracking on the treadmill.

(C) Cross-correlation coefficient between predicted and measured calcium signals per trial at a time lag of zero. The black line shows the median. The black dot marks the trial shown in (B). n: number of trials; N: number of flies.

(D) Predicted and measured calcium signals aligned to the transitions into and out of movement. Movement includes walking and grooming. Signals are baseline subtracted (mean from -0.5 to 0 s). Thin lines show animal means, thick lines show mean of means, shadings show standard error of the mean. n: number of transitions; N: number of flies.

(E) Example trial of two-photon calcium imaging of hook extension axons and behavior tracking without the treadmill.

(F) Same as (C) but for hook extension axons imaged without the treadmill.

(G) Same as (D) but for hook extension axons imaged without the treadmill.
(H) Example trial of two-photon calcium imaging of hook extension axons in the neuromere of the left front leg and behavior tracking on the platform.
(I) Same as (C) but for hook extension axons imaged on the platform. Active movements were excluded for the cross-correlation.
(J) Same as (D) but for hook extension axons imaged on the platform.
(K) Median predicted and measured calcium signals during active and passive movement bouts on the platform. Bouts are ≥0.5 s in duration. Distributions show kernel density estimations. n: number of movement bouts; N: number of flies.

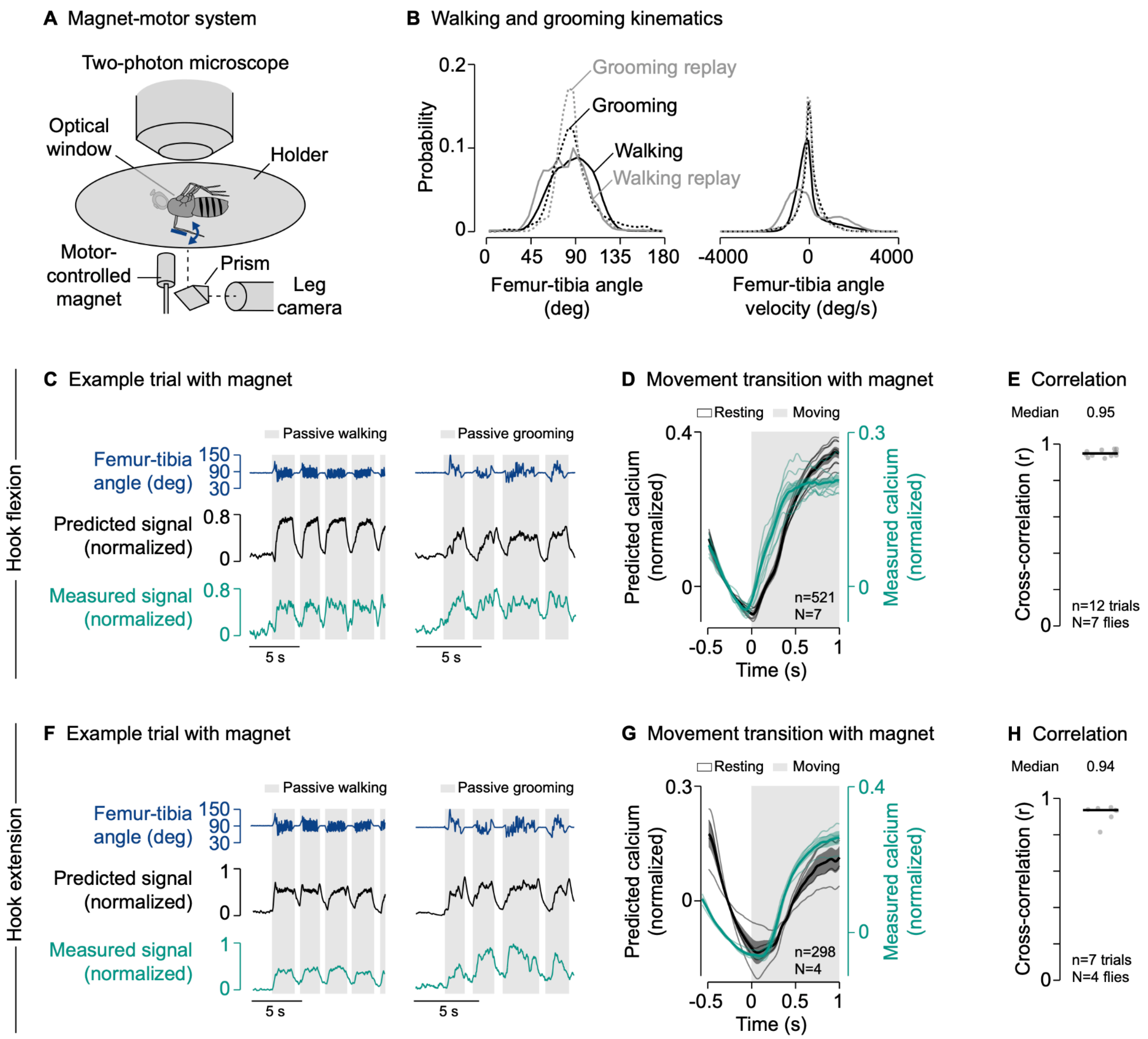


**Figure S5. Related to Figure 3**

(A) Experimental setup for two-photon calcium imaging from neurons in the neuromere of the left front leg and leg tracking in a tethered fly. All joints except for the femur-tibia joint of a front leg are fixated. The femur-tibia joint is passively moved via a motor-controlled magnet.

(B) Probability distributions of walking and grooming kinematics recorded in the hook flexion neuron dataset and the walking and grooming kinematics used for passive replay with the setup shown in (A).

(C) Example trial of two-photon calcium imaging of hook flexion axons in the neuromere of the left front leg and behavior tracking with the magnet-motor system.

(D) Predicted and measured calcium signals aligned to the transition into passive movement. Movement includes passive walking and passive grooming. Signals are baseline subtracted (mean from -0.5 to 0 s). Thin lines show animal means, thick lines show mean of means, shadings show standard error of the mean. n: number of transitions; N: number of flies.

(E) Cross-correlation coefficient between predicted and measured calcium signals per trial at a time lag of zero. The black line shows the median. Trials are either walking or grooming replay. n: number of trials; N: number of flies.

(F) Same as (C) but for hook extension axons.

(G) Same as (D) but for hook extension axons.

(H) Same as (E) but for hook extension axons.

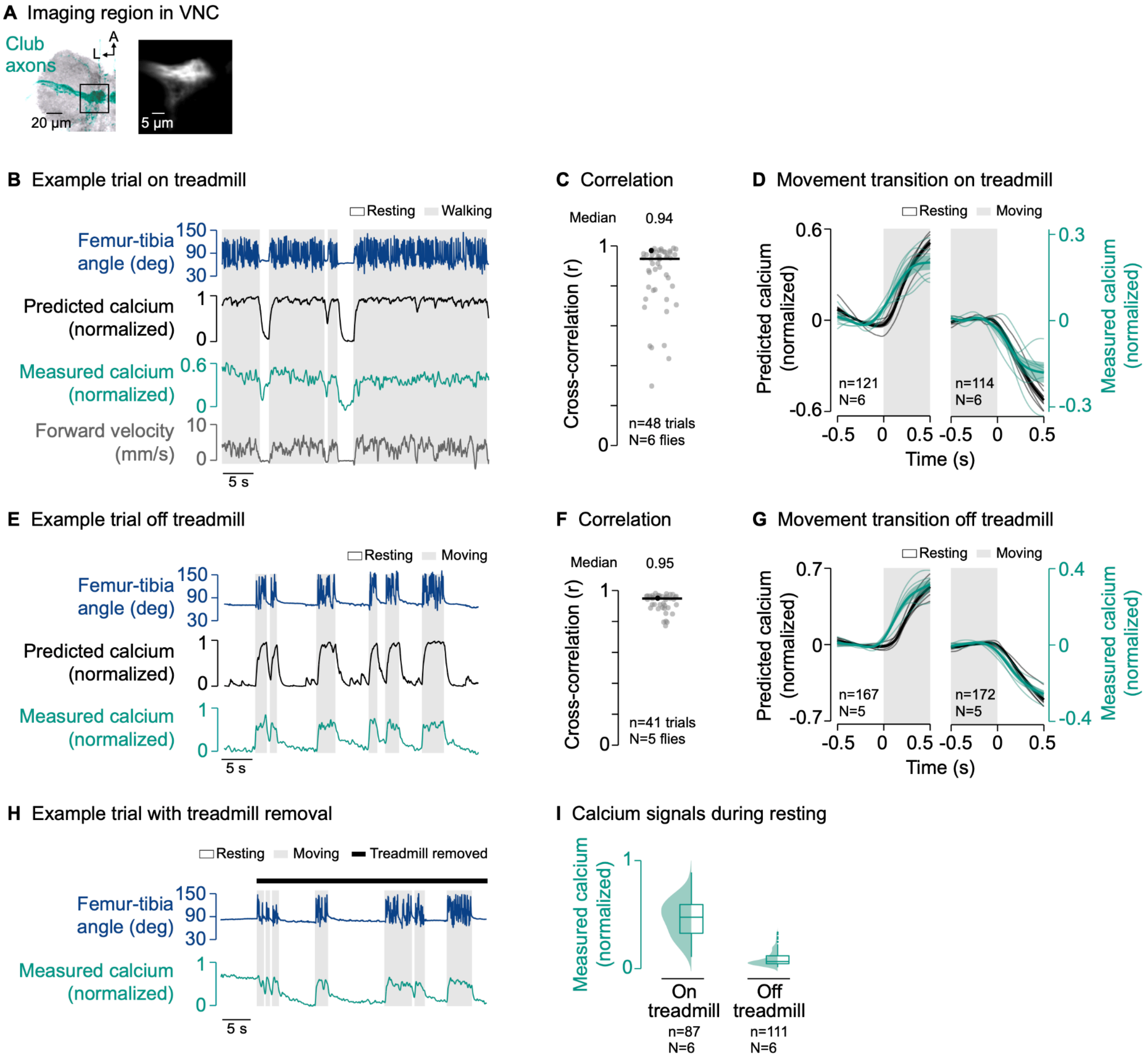


**Figure S6. Related to Figure 3**

(A) Left: Confocal image of club axons in the neuromere of the left front leg. The black box indicates the imaging region. Green: GFP; gray: neuropil stain (nc82). A: anterior; L: lateral. Right: Mean tdTomato signal within the imaging region during an example trial.

(B) Example trial of two-photon calcium imaging of club axons in the neuromere of the left front leg and behavior tracking on the treadmill.

(C) Cross-correlation coefficient between predicted and measured calcium signals per trial at a time lag of zero. The black line shows the median. The black dot marks the trial shown in (B). n: number of trials; N: number of flies.

(D) Predicted and measured calcium signals aligned to the transitions into and out of movement. Movement includes walking and grooming. Signals are baseline subtracted (mean from -0.5 to 0 s). Thin lines show animal means, thick lines show mean of means, shadings show standard error of the mean. n: number of transitions; N: number of flies.

(E) Example trial of two-photon calcium imaging of club axons in the neuromere of the left front leg and behavior tracking without the treadmill.

(F) Same as (C) but for club axons imaged without the treadmill.

(G) Same as (D) but for club axons imaged without the treadmill.

(H) Example trial of two-photon calcium imaging of club axons in the neuromere of the left front leg and behavior tracking. In this example, the treadmill was lowered (“removed”) about 5 s into the trial so the fly’s legs could not touch it.

(I) Median calcium signals during resting bouts on and off the treadmill. Bouts are ≥1 s in duration. Distributions show kernel density estimations. n: number of movement bouts; N: number of flies.

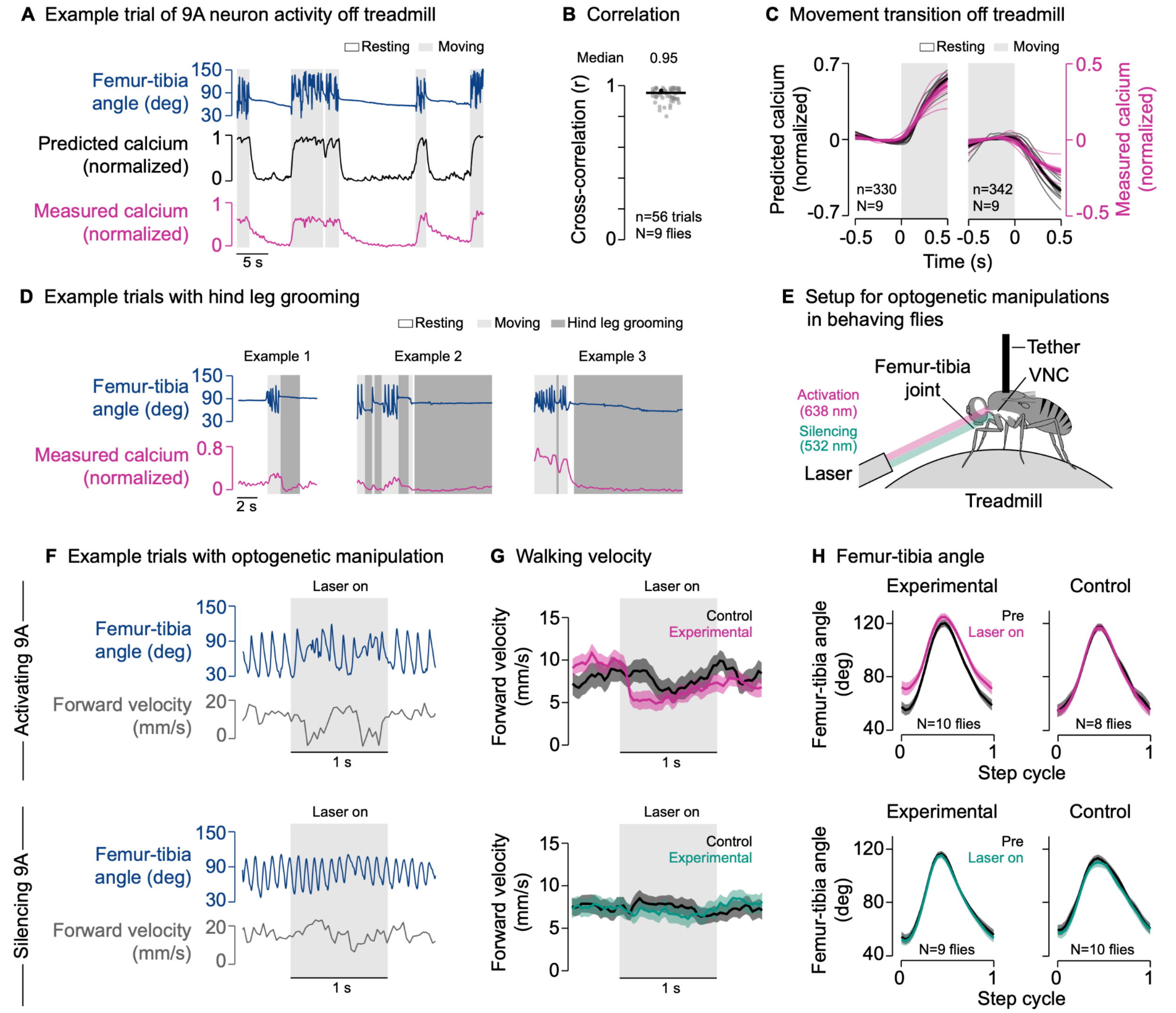


**Figure S7. Related to Figure 4**

(A) Example trial of two-photon calcium imaging of 9A neurons in the neuromere of the left front leg and behavior tracking without the treadmill.

(B) Cross-correlation coefficient between predicted and measured calcium signals per trial at a time lag of zero. The black line shows the median. The black dot marks the trial shown in (A). n: number of trials; N: number of flies.

(C) Predicted and measured calcium signals aligned to the transition into and out of movement. Signals are baseline subtracted (mean from -0.5 to 0 s). Thin lines show animal means, thick lines show mean of means, shadings show standard error of the mean. n: number of transitions; N: number of flies.

(D) Examples of two-photon calcium imaging of 9A neurons in the neuromere of the left front leg during hind leg grooming.

(E) Experimental setup for optogenetic manipulations of neurons in the neuromere of the left front leg of tethered flies walking on a treadmill. A red laser is used for optogenetic activation (via CsChrimson), a green laser is used for optogenetic silencing (via GtACR1). The fly's wings are trimmed.

(F) Example trials of optogenetic activation (top) and silencing (bottom) of 9A neurons in experimental flies walking on the treadmill.

(G) Average forward velocity of experimental and control flies during the 2 s trials. Thick lines show means of animal means, shadings show 95% confidence intervals of animal means. In experimental flies, 9A neurons are activated (top) or silenced (bottom). Control flies have the same genetic background but lack GAL4 expression.

(H) Femur-tibia angle of the left front leg of experimental and control flies normalized in time to the step cycle. Thick lines show means of animal means, shadings show 95% confidence intervals of animal means. Pre indicates steps recorded in the 0.5 s prior to laser onset. Laser on indicates steps recorded during the 1 s laser stimulation. In experimental flies, 9A neurons are activated (top) or silenced (bottom). Control flies have the same genetic background but lack GAL4 expression.

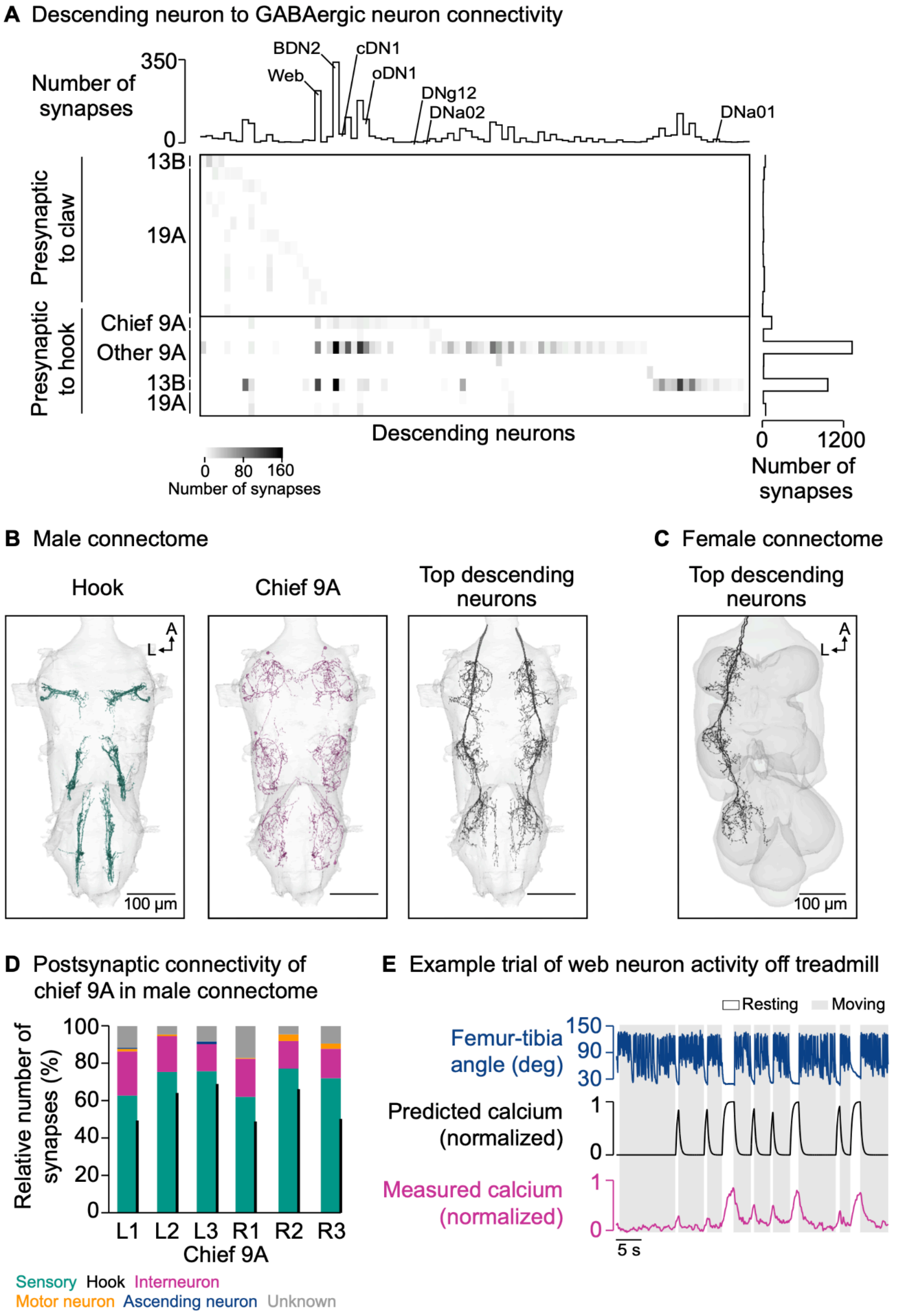


**Figure S8. Related to Figure 5**

(A) Connectivity of descending neurons with GABAergic neurons presynaptic to claw and hook axons. The grayscale heatmap indicates the number of synapses between neurons (connection strength). BDN2, cDN1, and oDN1 promote walking (Sapkal et al. 2023). DNa01 and DNa02 promote turning (Yang et al. 2023). DNg12 promotes grooming (Guo et al. 2022).

(B) Hook axons, chief 9A neurons presynaptic to hook axons, and the top two descending neurons presynaptic to the chief 9A neurons in the male VNC connectome (MANC). A: anterior; L: lateral.

(C) Top two descending neurons presynaptic to the chief 9A neuron in the female VNC connectome (FANC). A: anterior; L: lateral.

(D) Outputs of chief 9A neurons onto different neuron types (MANC connectome). Black bars indicate output onto hook axons. L: left side of the VNC; R: right side of the VNC; 1: front leg neuromere; 2: middle leg neuromere; 3: hind leg neuromere.

(E) Example trial of two-photon calcium imaging of the web neuron in the neuromere of the left front leg and behavior tracking without the treadmill.

**Video S1**
Neurons of interest in the FANC connectome.

**Video S2**
Example trials of two-photon calcium imaging of claw axons and behavior tracking on the treadmill and without the treadmill. Videos are sped up 2x. The measured calcium signal is based on the ratio of GCaMP to tdTomato. For this video, GCaMP images were low-pass filtered using a moving average filter with a time window of 0.2 s. tdTomato images are not shown.

**Video S3**
Example trials of two-photon calcium imaging of hook flexion axons and behavior tracking on the treadmill, without the treadmill, and on the platform. Videos are sped up 2x. The measured calcium signal is based on the ratio of GCaMP to tdTomato. For this video, GCaMP images were low-pass filtered using a moving average filter with a time window of 0.2 s. tdTomato images are not shown.

**Video S4**
Example trials of two-photon calcium imaging of club axons and behavior tracking on the treadmill, without the treadmill, and when the treadmill is removed during the trial. Videos are sped up 2x. The measured calcium signal is based on the ratio of GCaMP to tdTomato. For this video, GCaMP images were low-pass filtered using a moving average filter with a time window of 0.2 s. tdTomato images are not shown.

**Video S5**
Example trials of two-photon calcium imaging of 9A axons and behavior tracking on the treadmill, without the treadmill, and on the platform. Videos are sped up 2x. The measured calcium signal is based on the ratio of GCaMP to tdTomato. For this video, GCaMP images were low-pass filtered using a moving average filter with a time window of 0.2 s. tdTomato images are not shown.

**Video S6**
Example trials of two-photon calcium imaging of web axons and behavior tracking on the treadmill and without the treadmill. Videos are sped up 2x. The measured calcium signal is based on the ratio of GCaMP to tdTomato. For this video, GCaMP images were low-pass filtered using a moving average filter with a time window of 0.2 s. tdTomato images are not shown.

### Table S1. Neurons of interest in the FANC connectome

| Name in this study | Description | Neurotransmitter | ID |
|---|---|---|---|
| Claw | Sensory axon in T1L | ACH | 648518346500641203 |
| Claw | Sensory axon in T1L | ACH | 648518346498453780 |
| Claw | Sensory axon in T1L | ACH | 648518346499722309 |
| Claw | Sensory axon in T1L | ACH | 648518346483783343 |
| Claw | Sensory axon in T1L | ACH | 648518346496335735 |
| Claw | Sensory axon in T1L | ACH | 648518346486653999 |
| Claw | Sensory axon in T1L | ACH | 648518346474593218 |
| Claw | Sensory axon in T1L | ACH | 648518346500681907 |
| Claw | Sensory axon in T1L | ACH | 648518346477981653 |
| Claw | Sensory axon in T1L | ACH | 648518346484687181 |
| Claw | Sensory axon in T1L | ACH | 648518346489569900 |
| Claw | Sensory axon in T1L | ACH | 648518346487912272 |
| Claw | Sensory axon in T1L | ACH | 648518346508752447 |
| Claw | Sensory axon in T1L | ACH | 648518346514361543 |
| Claw | Sensory axon in T1L | ACH | 648518346481759933 |
| Claw | Sensory axon in T1L | ACH | 648518346490596093 |
| Claw | Sensory axon in T1L | ACH | 648518346488815453 |
| Hook | Sensory axon in T1L | ACH | 648518346480125925 |
| Hook | Sensory axon in T1L | ACH | 648518346476657526 |
| Hook | Sensory axon in T1L | ACH | 648518346481857725 |
| Hook | Sensory axon in T1L | ACH | 648518346514448583 |
| Hook | Sensory axon in T1L | ACH | 648518346480666625 |
| Hook | Sensory axon in T1L | ACH | 648518346509569667 |
| Hook | Sensory axon in T1L | ACH | 648518346507233352 |
| Hook | Sensory axon in T1L | ACH | 648518346494933426 |
| Hook | Sensory axon in T1L | ACH | 648518346494932914 |
| Hook | Sensory axon in T1L | ACH | 648518346494264434 |
| Hook | Sensory axon in T1L | ACH | 648518346481856445 |
| Hook | Sensory axon in T1L | ACH | 648518346489572037 |
| Hook | Sensory axon in T1L | ACH | 648518346501288024 |
| Hook | Sensory axon in T1L | ACH | 648518346477034696 |
| Hook | Sensory axon in T1L | ACH | 648518346494933170 |
| Hook | Sensory axon in T1L | ACH | 648518346486753761 |
| Hook | Sensory axon in T1L | ACH | 648518346496729980 |

| | | | |
|---|---|---|---|
| Hook | Sensory axon in T1L | ACH | 648518346494071307 |
| 13B | Interneuron in T1L presynaptic to claw | GABA | 648518346484847261 |
| 19A | Interneuron in T1L presynaptic to claw | GABA | 648518346531401754 |
| 19A | Interneuron in T1L presynaptic to claw | GABA | 648518346472653065 |
| 19A | Interneuron in T1L presynaptic to claw | GABA | 648518346488555913 |
| 19A | Interneuron in T1L presynaptic to claw | GABA | 648518346498429489 |
| 19A | Interneuron in T1L presynaptic to claw | GABA | 648518346502916595 |
| 19A | Interneuron in T1L presynaptic to claw | GABA | 648518346488659278 |
| 19A | Interneuron in T1L presynaptic to claw | GABA | 648518346521506809 |
| 19A | Interneuron in T1L presynaptic to claw | GABA | 648518346518741215 |
| 19A | Interneuron in T1L presynaptic to claw | GABA | 648518346465091957 |
| 19A | Interneuron in T1L presynaptic to claw | GABA | 648518346494217863 |
| 19A | Interneuron in T1L presynaptic to claw | GABA | 648518346499593822 |
| 19A | Interneuron in T1L presynaptic to claw | GABA | 648518346501344355 |
| 19A | Interneuron in T1L presynaptic to claw | GABA | 648518346491096865 |
| 3A | Interneuron in T1L presynaptic to claw | ACH | 648518346499860364 |
| Chief 9A | Interneuron in T1L presynaptic to hook | GABA | 648518346496946148 |
| 9A | Interneuron in T1L presynaptic to hook | GABA | 648518346479847574 |
| 9A | Interneuron in T1L presynaptic to hook | GABA | 648518346479837078 |
| 9A | Interneuron in T1L presynaptic to hook | GABA | 648518346498002535 |
| 9A | Interneuron in T1L presynaptic to hook | GABA | 648518346479879156 |
| 9A | Interneuron in T1L presynaptic to hook | GABA | 648518346467364359 |
| 9A | Interneuron in T1L presynaptic to hook | GABA | 648518346486716621 |
| 13B | Interneuron in T1L presynaptic to hook | GABA | 648518346502572199 |
| 19A | Interneuron in T1L presynaptic to hook | GABA | 648518346479427282 |
| 19A | Interneuron in T1L presynaptic to hook | GABA | 648518346494008718 |
| 8A | Interneuron in T1L presynaptic to hook | GLUT | 648518346488868849 |
| 1A | Interneuron in T1L presynaptic to hook | ACH | 648518346488991501 |
| 8B | Intersegmental ascending neuron presynaptic to hook | ACH | 648518346489674348 |
| 8B | Intersegmental ascending neuron presynaptic to hook | ACH | 648518346479174095 |
| 18B | Intersegmental ascending neuron presynaptic to hook | ACH | 648518346490380810 |
| 18B | Intersegmental ascending neuron presynaptic to hook | ACH | 648518346473080420 |
| 22A | Interneuron in T1L presynaptic to hook | ACH | 648518346518548566 |
| Hook | Sensory axon presynaptic to hook | ACH | 648518346480125925 |
| Hook | Sensory axon presynaptic to hook | ACH | 648518346494933170 |

| | | | |
|---|---|---|---|
| Hook | Sensory axon presynaptic to hook | ACH | 648518346481857725 |
| Hook | Sensory axon presynaptic to hook | ACH | 648518346476657526 |
| Hook | Sensory axon presynaptic to hook | ACH | 648518346494933426 |
| Hook | Sensory axon presynaptic to hook | ACH | 648518346490237960 |
| Hook | Sensory axon presynaptic to hook | ACH | 648518346496671612 |
| Hook | Sensory axon presynaptic to hook | ACH | 648518346514448583 |
| Hook | Sensory axon presynaptic to hook | ACH | 648518346501288024 |
| Hair plate | Sensory axon presynaptic to hook | ACH | 648518346499992716 |
| Unknown | Neurite in T1L presynaptic to hook | N/A | 648518346494610152 |
| Web | Descending neuron presynaptic to chief 9A | N/A | 648518346478690132 |
| BDN2 | Descending neuron presynaptic to chief 9A | N/A | 648518346459693060 |
| Descending neuron | Descending neuron presynaptic to chief 9A | N/A | 648518346476980936 |
| Descending neuron | Descending neuron presynaptic to chief 9A | N/A | 648518346459681796 |
| oDN1 | Descending neuron presynaptic to chief 9A | N/A | 648518346504806022 |
| Descending neuron | Descending neuron presynaptic to chief 9A | N/A | 648518346485765060 |
| Descending neuron | Descending neuron presynaptic to chief 9A | N/A | 648518346479290513 |
| DNa02 | Descending neuron presynaptic to chief 9A | N/A | 648518346478550356 |
| Descending neuron | Descending neuron presynaptic to chief 9A | N/A | 648518346475284321 |
| Descending neuron | Descending neuron presynaptic to chief 9A | N/A | 648518346475392384 |
| Descending neuron | Descending neuron presynaptic to chief 9A | N/A | 648518346484448003 |
| DNg12 | Descending neuron presynaptic to chief 9A | N/A | 648518346498347057 |
| Descending neuron | Descending neuron presynaptic to chief 9A | N/A | 648518346496855192 |
| cDN1 | Descending neuron presynaptic to chief 9A | N/A | 648518346520017233 |
| Descending neuron | Descending neuron presynaptic to chief 9A | N/A | 648518346494165305 |
| Descending neuron | Descending neuron presynaptic to chief 9A | N/A | 648518346487779095 |
| Descending neuron | Descending neuron presynaptic to chief 9A | N/A | 648518346490241884 |
| Descending neuron | Descending neuron presynaptic to chief 9A | N/A | 648518346477810925 |
| Descending neuron | Descending neuron presynaptic to chief 9A | N/A | 648518346496600109 |

IDs are from FANC CAVE materialization version 840, timestamp 2024-01-17T08:10:01.179472.

### Table S2. Neurons of interest in the MANC connectome

| Name in this study | Description | Neurotransmitter | ID or group name |
|---|---|---|---|
| Hooks | Sensory axons from hook neurons in all leg neuropils | ACH | SNpp38 |
| Chief 9A | Interneuron in T1L presynaptic to hook | GABA | 100513 |
| Chief 9A | Interneuron in T2L presynaptic to hook | GABA | 13157 |
| Chief 9A | Interneuron in T3L presynaptic to hook | GABA | 14517 |
| Chief 9A | Interneuron in T1R presynaptic to hook | GABA | 165560 |
| Chief 9A | Interneuron in T2R presynaptic to hook | GABA | 12443 |
| Chief 9A | Interneuron in T3R presynaptic to hook | GABA | 12804 |
| Web | Descending neuron presynaptic to chief 9A in T1L, T2L, T3L | GABA | 10107 (DNxl041) |
| Web | Descending neuron presynaptic to chief 9A in T1R, T2R, T3R | GABA | 10103 (DNxl041) |
| Descending neuron | Descending neuron presynaptic to chief 9A in T1L, T2L, T3L | ACH | 10093 (DNxl058) |
| Descending neuron | Descending neuron presynaptic to chief 9A in T1R, T2R, T3R | ACH | 10339 (DNxl058) |

IDs are from MANC version 1.0.

### Table S3. Neurons of interest in the FlyWire connectome

| Name in this study | Description | Neurotransmitter | ID or group name |
| --- | --- | --- | --- |
| Web | Descending neuron presynaptic to chief 9A in T1L, T2L, T3L | GABA | 720575940636656632 (DNg74_b) |
| Web | Descending neuron presynaptic to chief 9A in T1R, T2R, T3R | GABA | 720575940627087646 (DNg74_b) |

IDs are from FlyWire public release version 783.
